# Predicting hypersensitivity and comorbid depressive-like behavior in late stages of joint disease using early weight bearing deficit

**DOI:** 10.1101/2023.11.29.569246

**Authors:** Sara Hestehave, Roxana Florea, Alexander J.H. Fedorec, Maria Jevic, Lucile Mercy, Annia Wright, Oakley B. Morgan, Laurence A. Brown, Stuart N. Peirson, Sandrine M. Géranton

## Abstract

Chronic pain is a hallmark of joint diseases and is often accompanied by negative affective symptoms such as low mood, anxiety and memory dysfunction. Whether these may be the results of the more obvious sensory and functional symptoms of joint pain is poorly understood and this likely contributes to the difficulty in adequately managing this condition. Here, we have used two mouse models to address this lack of knowledge. Using a model of ankle inflammation and a model of knee osteoarthritis, we found that these models of joint pain induced weight bearing deficits of different magnitude but relatively similar mechanical allodynia that lasted at least 3 months. However, the models were accompanied by very different affective outcomes, as only the model of knee osteoarthritis, that led to significant early changes in activity and sleep patterns, was accompanied by an increase in negative affective behaviors, including cognitive impairments and depressive-like behavior. The models also had different molecular profiles at both spinal and hippocampal levels. Importantly, the functional outcomes measured in the early stages of the disease stage strongly correlated with sensory and emotional profiles at 3 months, suggesting that early functional measures may be used as predictors of the long-term symptoms associated with persistent joint pain. In conclusion, the predictive value of early measures of functional impact of joint disease could prove useful in the clinics for adapted therapeutic approaches for the prevention of emotional comorbidities and better pain management for patients with joint pain.

## Introduction

Patients with chronic pain often experience negative affective symptoms such as low mood, anxiety and disturbed sleep, that altogether significantly impact patients’ well-being. While the sensory aspects of persistent pain have been extensively studied in animal models, associated mood-related disorders, that are a crucial component of the human experience of chronic pain, are not often enough assessed in pre-clinical research, and even less so in joint-pain models. As a result, the relationship between affective symptoms and persistent hypersensitive states remains poorly understood. Joint disease is one of the most widespread chronic diseases associated with persistent pain. Importantly, the prevalence of clinically significant anxiety and/or depression in patients with joint pain is at least 41% [5]. While the pain severity is often not related to the extent of the joint damage [4,35], it was found to positively correlate with anxiety and depression [46]. However, little is known about the relationship between the early signs of joint disease, such as gait changes and hypersensitivity, with the long-term affective outcomes. A better understanding of how these may be connected could support better pain management and help with the prevention of significant comorbidities by early intervention.

Here, we have used two well described models of joint pain with fast onset of mechanical hypersensitivity for which we knew very little about potential comorbid affective behaviors. We used the model of Complete Freund’s Adjuvant (CFA)-induced ankle joint inflammation [20,34], and the monoiodoacetate (MIA) model of knee osteoarthritis [39,40,43,47]. Intra-articular injection of CFA leads to the infiltration of inflammatory cells and synovial hypertrophy and generates robust and long-lasting pain-like behaviours, and much is known about the central mechanisms involved in the development of this pain state [20]. This model has been associated with anxiety- and depression-like behaviour in previous studies, but the time course of development of these behaviours remains unclear [30]. The MIA-model induces rapid disease progression in the knee, as MIA inhibits the glycolytic pathway causing rapid and widespread chondrocyte death, extensive neovascularization, subchondral bone necrosis and collapse, as well as profound and prolonged inflammation [17]. This model induces long-lasting mechanical hypersensitivity and is responsive to conventional pain-relieving therapies [15,43]. Importantly, intra-articular injection of MIA produces histological alterations similar to those found in clinical histopathology from arthritis patients [22,28].

Using both models simultaneously with control mice only exposed to brief anaesthesia, we monitored animals with a range of weight bearing deficits and mechanical hypersensitivity, allowing us to look at the correlations between functional, sensory, affective and molecular outcomes in the early and late stages of the disease states. We report that both models had relatively similar sensory profiles across the 3 months of monitoring of the animals’ behaviour, but that there were significant differences in weight bearing deficit and emotional behaviors that only developed after MIA. Importantly, depression-like behaviour that developed 3 months after the onset of the joint pain correlated with weight bearing deficit, not only measured at the late stage but also measured at the early stage of the disease state. These results suggest that early functional outcomes of joint disease could be used to predict the likelihood to develop emotional comorbidities.

## Methods and materials

### Animals and housing

This study focused on male mice, as its aim was to explore differences between pain models in terms of sensory, functional and emotional outcomes. We therefore decided to strengthen our statistical power towards treatment outcome (i.e. CFA *vs* MIA) which necessitate high n numbers when looking at emotional parameters, as suggested by our initial power calculations. Moreover, the oestrus cycle is known to affect locomotor activity patterns in females, which would have meant much longer monitoring and larger sample sizes needed for our activity and sleeping patterns studies. For all experiments, male mice (C57Bl/6J from Charles River, UK) arrived at our facility at 8 weeks of age, and were left to acclimatize for at least 7 days before experiments started All animals were kept in groups of 4-5 in a temperature-controlled (20±1°C) environment in Individual Ventilated Cages-cages; cardboard tunnels for shelter; light-dark cycle of 12 hours (gradual lights on between 7-8a.m. and off between 7-8p.m); and ad libitum provision of food and water. Only exception to these housing-conditions was during the sleep/activity-recordings, where animals were single-housed without tunnels for shelter. All experiments were carried out under the Home Office License P8F6ECC28, and all efforts were made to minimize animal suffering and to reduce the number of animals used (UK Animal Act, 1986).

### Study design

The study was divided in cohorts/studies including animals from all experimental groups (CFA vs MIA vs control, n = 4-7). In order to not stress the mice from over-testing, and thereby confounding the results, each experiment was designed to test only a selection of parameters. Experiments were also terminated at different time points in order to collect tissue at the appropriate time point for various molecular markers. Therefore, group-sizes were variable throughout the study, but always distributed equally across treatment. Group-sizes are presented as a range throughout the manuscript.

### Experimental procedures

#### Induction of injury, (CFA)

Induction of the Complete Freund’s Adjuvant model of tibio-tarsal joint inflammation (CFA) was performed similar to what was previously reported by our group [33]. Anesthesia was induced similarly to above, but the animal was placed in lateral recumbency on the right side, for fixation of the left ankle joint. Using a precision Hamilton-syringe, a 25G needle entered the ankle joint from the lateral posterior position, with the ankle kept in plantar flexion to open the joint, and 5µl of CFA (Complete Freund’s Adjuvant, Sigma) was injected. Control animals were only exposed to anesthesia.

#### Induction of injury, (MIA)

Induction of the Monoiodoacetate Arthritis Model to the knee joint (MIA) was performed similar to previously reported [43]. On the day of injection, the monoiodoacetate solution was freshly prepared in sterile saline for injection of 1mg Sodium Iodoacetate (≥98%, Sigma, I2512-25G,) in 10µl saline (9% NaCl). Animals were anaesthetized in an induction chamber using Isoflurane 2.5% mixed in O_2_ at a flowrate of 1.5 L/min, and maintained via facemask at 2.0% during the injection. Anaesthetic depth was confirmed by lack of withdrawal reflex to a pinch to the tail. The animal was then placed on the back in dorsal recumbency, and the fur was shaved in the area around the knee on the left hindleg. The knee was stabilized and fixed in a slightly bend position and the patellar tendon was visualized as a white line below the skin. The injection of 10µl using a 30G insulin-syringe (BD Micro-Fine Plus Demi, 0.3ml (30G) 8mm) was made intraarticularly in the joint-space by applying it perpendicularly through the tendon just below the patella, and with as minimal movement as possible. Note that the compound is very toxic, and that even small volumes delivered subcutaneously outside of the joint-space, or potential damage to the popliteal artery in the knee, may cause intoxication of the animal. Great care must be taken to secure a controlled injection and to secure accurate volume to not risk leakage of the fluid from the joint space.

### Behavioural testing

Behavioural testing was always performed in randomized order and by the same female experimenter. Animals were always allowed at least 30min of habituation to the testing room prior to behavioural testing. Unless otherwise specified behavioral tests were always performed between 8am and 2pm.

#### Mechanical allodynia (VF)

Low intensity mechanical sensitivity assessment was similar to previously reported by our group [34], using a series of calibrated von Frey monofilaments (0.02; 0.04; 0.07; 0.16; 0.4; 0.6; 1.0) (Ugo Basile SRL, Italy). Animals were placed in Plexiglas chambers, located on an elevated wire grid, and allowed to habituate to the testing environment for at least 60min prior testing. Once the animals were calm, the plantar surface of the paw was stimulated, starting with a 0.6g filament applied with uniform pressure for 5seconds. A brisk withdrawal, stretching or licking of toes, was considered as a positive response, whereupon the next lower-force filament was applied. In the absence of a positive response, the next higher-force filament was applied in the next test. After the first change in response-pattern, suggesting the threshold, additional 4 stimulations were applied; applying the next higher-force filament when no response, and lower-force filament following positive responses. The response pattern determined the constant, *k* [16], and the 50% response threshold was determined using the following equation; 50% threshold (g) = 10^log(last^ ^filament)+k*0.3^.

#### Affective-motivational behaviour (AF)

In addition to assessing pure reflexive sensory threshold to mechanical stimulation (VF), we also adopted and modified the protocol described by [13,32] to assess the affective responses displayed after stimulation with three selected filaments (low; 0.04g, medium; 0.16g, high; 1.0g). The assessment was always performed following the VF-assessment, while the animals were still in the Plexiglas chamber, and following a period of rest. Starting with the low-intensity filament, each filament was applied once for 1sec, and the duration of affective response was recorded as the amount of time the animal showed conscious attending behavior to the stimulated paw by licking, biting, lifting, looking at or guarding the paw, within 30 seconds after application of the filament. The animal was left to rest for at least 10min before the next higher filament force was applied.

#### Cold allodynia, (ADT)

While the animals were still in the plexiglas chambers following von Frey measurements, cold allodynia was assessed using application of a drop of acetone (Acetone Drop Test, ADT) to the plantar surface of the paw, using a plastic syringe without mechanically touching the skin. Following application, the duration of the response was then recorded, with a maximum of 30 seconds. A positive response was considered as flinching, licking, looking at or withdrawing the paw. The application and assessment was performed 2-3 times on each paw for each animal with 5-10 minutes between each application, and the average of the measurements was calculated.

#### Functional impairment - Weight bearing (WB)

To assess the functional impact of the injury the static weight bearing distribution was assessed similarly to previously described [29]. Hindlimb weight bearing was measured using a Bioseb Incapacitance Test (Bioseb) which measures the weight distribution across the two hindlimbs of a stationary animal. Animals were habituated and trained to become comfortable with the testing paradigm during short sessions for 5 days prior to induction of the model. Three readings were collected and averaged for each animal, and the weight borne by the ipsilateral limb was expressed as a percentage of the weight borne across both hindlimbs (WB-% = (weight borne on the injured leg/weight borne on both legs) * 100%).

#### Catwalk gait analysis

Analysis of voluntary movement and gait pattern was performed using the Catwalk® XT 10.0 system (Noldus Information Technology) [30], and based on our previous experience [23]. For optimal contrast for the recording, testing was conducted in a dark room. Briefly, green light was internally reflected into a glass plate, on which an enclosed corridor was fixed, with red backlight above the corridor. A video-camera was mounted underneath the setup and recorded the paw prints being lit up by the green light when paws were in contact with the glass plate as the animal walked along the corridor. A run was regarded as compliant when the animal entered in one end of the corridor, and moved fluently across the plate towards the other end of the corridor, with a running duration below 12 seconds and a maximum variation below 75%. Three compliant runs were recorded for each animal, with no previous training/habituation or food-deprivation. Following the recording, compliant runs were classified and cleaned/corrected for potential miscellaneous prints. For outcome-measures like swing-time ratio, contact area and single stance, the data was converted into a ratio between ipsi- and contra-lateral hind-limbs. The term “print position” refers to the distance between the position of the hindpaw, and the previously placed front paw. No prior habituation to the equipment or training was needed for the animals to complete the desired compliant runs.

#### Sucrose Preference Test (SPT)

In order to assess depressive-like behaviour, the sucrose preference test was included as a measure of anhedonia [49], At least 5 days before SPT each home-cage was fitted with two water bottles, in order to have the mice accustomed to drinking from both bottles / sides of the cage. Two days before the SPT test, one of the water-bottles was filled with 1% sucrose solution for approximately 24h, with the two bottles changing side half way, to allow the mice time to discover the sweet solution, and learn that it may be presented in both sides. No sucrose was provided the last 24 hours before the actual test. For the SPT test, all animals were individually housed in clean cages similar to their home-cage environment for 12h overnight (7pm-7am), provided similar enrichment, nesting material and food on the same IVC-rack as their home-cage, and were given free access to two pre-weighed bottles containing normal drinking water or 1% sucrose solution. In the morning all bottles were weighed and mice were placed back together with their cage-mates. This re-introduction released some fighting in some of the cages, and mice were given the following night to calm down before the test was repeated overnight with the sucrose bottle presented in the opposite side, to account for potential side-preference, as reported previously in rats [23]. No side-preference was detected in the current study, and the sucrose preference % was calculated on the total amount consumed across the two nights combined, using the following calculation; SPT% = (sucrose-solution consumed / (total fluid consumption)) * 100%. The sucrose-solution was always prepared fresh before provision at 1% in the normal drinking water. As recent meta-analysis suggests food- and water-deprivation to confound the outcome of this assay [8] no water- or food-deprivation was employed in this study.

#### Elevated Plus Maze (EPM)

To assess anxiety-like behaviour an EPM was used as described previously [23,24], but for these experiments adopted to mice rather than rats. The maze consisted of four arms (35*5 cm) arranged in a cross-like disposition, and 60cm above ground (Ugo Basile SRL, Italy). Two opposite arms were open, and the other two were equipped with 15 cm high walls on each side for enclosement. All were connected by a central 5*5 cm square. The animal was placed in the centre, for free exploration for 5 minutes. Recording was performed by use of a camera placed above the maze, and movement between zones was tracked using EthoVision XT14 (Noldus Information Technology).

#### Open Field Test (OFT)

To assess anxiety-like and locomotor activity, an OFT was used. The OFT was a circular open arena with grey plastic flooring and blue plastic sides (diameter, 36cm, height 32cm). The arena was evenly illuminated by lighting placed above the arena. A video camera was positioned directly above the arena and connected to a computer performing live-tracking and recording of the behavior using Ethovision XT14. The animal was placed in the middle of the arena and allowed 5 minutes of freely exploration. The proportion of time spent in the centre vs by the edges/walls of the arena, was used as a marker for the anxiety-like behavior (outer zone; 6.7cm along the edge of the arena. Centre zone diameter; 22.6cm).

#### Novel Object Recognition (NOR)

The Novel Object Recognition test was performed similarly to previously described [9,37], with some modifications. Testing was carried out in the same circular arena as the OFT, and always on the following day, making the OFT also serve as habituation to the arena without objects. A video camera was positioned directly above the arena and connected to a computer performing live-tracking and recording of the behavior using Ethovision XT14, with 3-point tracking of nose-center-tail points. The familiar/similar objects were brown circular glass bottles (d; 7cm, h; 18cm), while the “novel” object was a translucent elliptical glass bottle (d; 9*5cm, h; 15cm) containing white sand for coloring. The animal was placed in the arena facing away from the objects, and first allowed 10minutes freely exploration and habituation to the arena including the two similar brown glass bottles (making these “familiar object” following the habituation phase), placed in two opposite quadrants of the arena. Following habituation, the animal was rejoined with its cage-mates. Three hours later, the animal was reintroduced to the arena for a 5min test, where one of the familiar objects had been replaced with the novel object, displaying a different color and shape, but same texture and sensation upon manipulating. The object replaced / side was alternated between test subjects and experimental groups to randomize for potential side-preferences. Upon completion, all recordings/trackings were corrected for appropriate nose/tail-tracking by the software, and exploration of an object was defined as the nose being within approximately 2cm of the objects. The proportion of time spent exploring the objects was assessed by calculating a percentage of time exploring the novel object, or for the habituation-phase; the object which was later replaced by the novel object. Using the following formula;

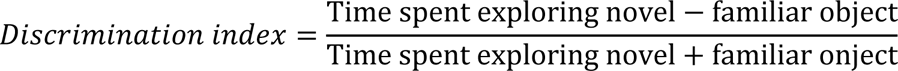

#### Joint circumference (JC)

Joint circumference was assessed to give an indication of the inflammatory development after the two injuries. It was measured on all left ankle and knee-joints in one cohort of animals, while the animal was already anaesthetized immediately before induction of the injury, and immediately before perfusion of the individual animal. The lateromedial (LM) and dorsoplantar (DP) diameters were measured using calipers, and the circumference was calculated using an approximation of the perimeter of an ellipse; 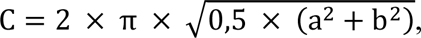 where a = the radius of DP and b = the radius of LM [6,7].

#### Sleep/activity-pattern

To measure undisturbed activity in the home-cages, we adopted the approach of Brown et al. [12] using non-invasive passive infrared motion sensors. Animals were single-housed with a 12h light:12h dark cycle. The cages, were open with wire tops (floor area: 500cm^2^) and a passive infrared (PIR) motion sensor was fitted above the cage. For accurate measurements of activity, the area below the food- and water-hopper was blocked off and tunnels / shelters were removed, allowing the animal to be in the sensors receptive field at all times. For these experiments, due to restriction on the time animals could remain single housed, animals were first injected and then placed directly into the recording cages, where they remained for a week of uninterrupted recording. Home cage mouse activity was tracked as in Brown et al. [12], with measurements taken every 10 seconds across multi-day periods. As before, sleep was defined as periods in which no activity was measured for 40 seconds or more [12]. Activity and sleep data were smoothed by calculating the mean in 10 minute bins as preliminary experiments demonstrated that this provided a balance between reducing measurement noise and maintaining time series features. Several summary statistics of circadian disruption [11] were calculated for individual animals across the 7 day period and, where appropriate, on each individual day: inter-daily stability, intra-daily variability, light-phase activity, dark-phase sleep, The Lomb-Scargle periodogram, similar to the chi-square periodogram [44].

#### Immunohistochemistry

For immunohistochemistry, mice were deeply anaesthetized with pentobarbital, weighed (BW), and perfused transcardially, first with heparinized saline (5000IU/mL), followed by freshly made 4% paraformaldehyde in 0.1M phosphate buffer (20mL pr adult mouse). Spinal cord and brain were dissected out and postfixed in the same paraformaldehyde solution for 2 hours, before being transferred to 30% sucrose solution in PB with 0.01% NaN3 at 4°C until cutting, at least 3 days after perfusion. The tissue was cut on a freezing microtome at 40μm thickness.

Lumbar spinal cord and hippocampal sections were rinsed in 0.1M PB and subsequently incubated at room temperature for 1h with 30% (v/v) normal goat serum (Invitrogen, Cat#31872) in 0.1M PB containing 10% Triton X-100. Sections were then incubated overnight at RT with one of the following primary antibodies: guinea pig anti-c-Fos (1:2000, Synaptic Systems, Cat#226004), rabbit anti-CGRP (1:5000, Millipore, AB5920), rabbit anti-IBA1 (1:500, Synaptic Systems, Cat#234008), rabbit anti-GFAP (1:4000, Dako, Z0334) or goat anti-DCX (1:500, Santa Cruz Biotechnology, F2916). Next day sections were washed three times in 0.1M PB and then incubated in darkness for 2h at RT with the respective secondary antibodies diluted in TTBS at 1:500. Lastly, after final washes, sections were mounted on gelatinized slides, coverslipped with Fluoromount Aqueous Mounting Medium (Sigma-Aldrich, F4680), and stored at 4°C.

#### Microscopy and quantification of immunofluorescence

Cell counting of c-Fos positive neurons (spinal cord sections from Lumbar L4 to L6) was performed directly under the microscope. For the other stains, images were collected using a Leica (Nussloch, Germany) DMR microscope connected to a Hamamatsu (C4742-95; Shizuoka, Japan) digital CCD camera and Volocity 6.3 Software. Intensity of CGRP immunoreactivity in superficial laminae LI-II as well as GFAP stain were analyzed using the ‘Mean grey value’ plugins in ImageJ. Background intensity of Laminae IV-V was subtracted from the positive signal in LI-II. Finally, IBA1 positive microglia in laminae I to III of the lumbar cord L4 to L6 and DCX positive cells in the dentate gyrus were manually counted from pictures using the counter feature in ImageJ.

#### Data and statistical analysis

All experiments were randomized and performed by a researcher unaware of the treatment groups. All statistical tests were performed in IBM SPSS Statistic Program (vers. 26) or GraphPad Prism (vers 9), and P<0.05 was considered statistically significant. Repeated Measures ANOVA was used when appropriate, but when some experiments/cohorts were terminated early due to the Covid-19 pandemic or tissue-collection, the data-sets affected were analyzed using Mixed-effects model to accommodate the missing data for the remaining period, while still including what had been obtained. Pearson r was used for correlation analysis. All details on statistical analysis, factors tested and significant outcomes, can be found in supplementary tables S1 and S2.

Weighted average was calculated as an Area Under the Curve for each individual animal tested, and the value was divided by the number of days in the experimental period, thereby giving a meaningful value to compare the overall differences between groups. For the majority of outcome measures, where we explored long term outcomes, we calculated a weighted average for two different periods defining the phases of the injury, “early” and “late”. The phases were defined as Early; from induction to around day 22, Late; from day 23 onwards (or the first data-collection/test following the day 23 test). The VF-data-set was log-transformed to ensure a normal distribution, as the von Frey hairs are distributed on an exponential scale. This was similar to previous studies in our group [34], and others [36].

We also used an approach of individual behavioural profiling, developed to differentiate between “affected” and “exposed but unaffected” animals [3,45]. Affected animals were defined as responding one standard deviation or more from the average performance of the control group. This calculation allowed to assess the percentage of affected animals within each treatment group.

## Results

### CFA and MIA induce functional deficit and persistent hypersensitivity that correlate for at least up to 3 months

The functional impact of joint disease was assessed using both static weight bearing and dynamic gait measures. Static bodyweight distribution has been used as a surrogate for the joint pain associated with weight-bearing and the static weight bearing assay has been the key outcome measure for unilateral joint-pain models for decades [10]. Here, we detected clear and prolonged weight bearing asymmetry for both the CFA and the MIA model (Fig.1A1), but with different characteristics. Because the early phase of the disease states seemed to indicate striking differences in weight bearing deficit between the 2 models, we decided to split the data between an early phase (day 0 to day 25) and a late phase (day 26 to day 90) as others have done before [52] (Fig.1A2). The MIA-model showed a strong weight bearing impairment in the early phase, after which it stabilized while the CFA-model produced a stable deficit across the time course of the observations (Fig 1A). Overall, the MIA-model caused a higher degree of weight-bearing asymmetry than the CFA-model (P<0.0001, for both the early and late phase). We next conducted a full gait analysis using the Catwalk which confirmed the biphasic nature of the weight bearing deficit in the MIA-model. Here, we present a selection of the most striking and significant parameters, all displayed as a ratio between left and right hind-leg, as for the static weight-bearing outcome. Overall, the Catwalk analysis detected gait-changes in the MIA model in the early phase on parameters related to Swing Time (Fig.1B), Duty Cycle, Stride Length, Single Stance (Fig.S1A-C) and also in Print Position (Fig.S1D), with clear correlations with the static weight bearing outcome at peak Day 8 and the early 3-week time point (Fig.S1E,F). In the late phase of the joint diseases, the dynamic gait-changes were relatively mild and the gait of injured animals assessed by the Catwalk analysis were very rarely different from the gait of control animals (Fig.1B, Fig.S1G).

**Figure 1.**
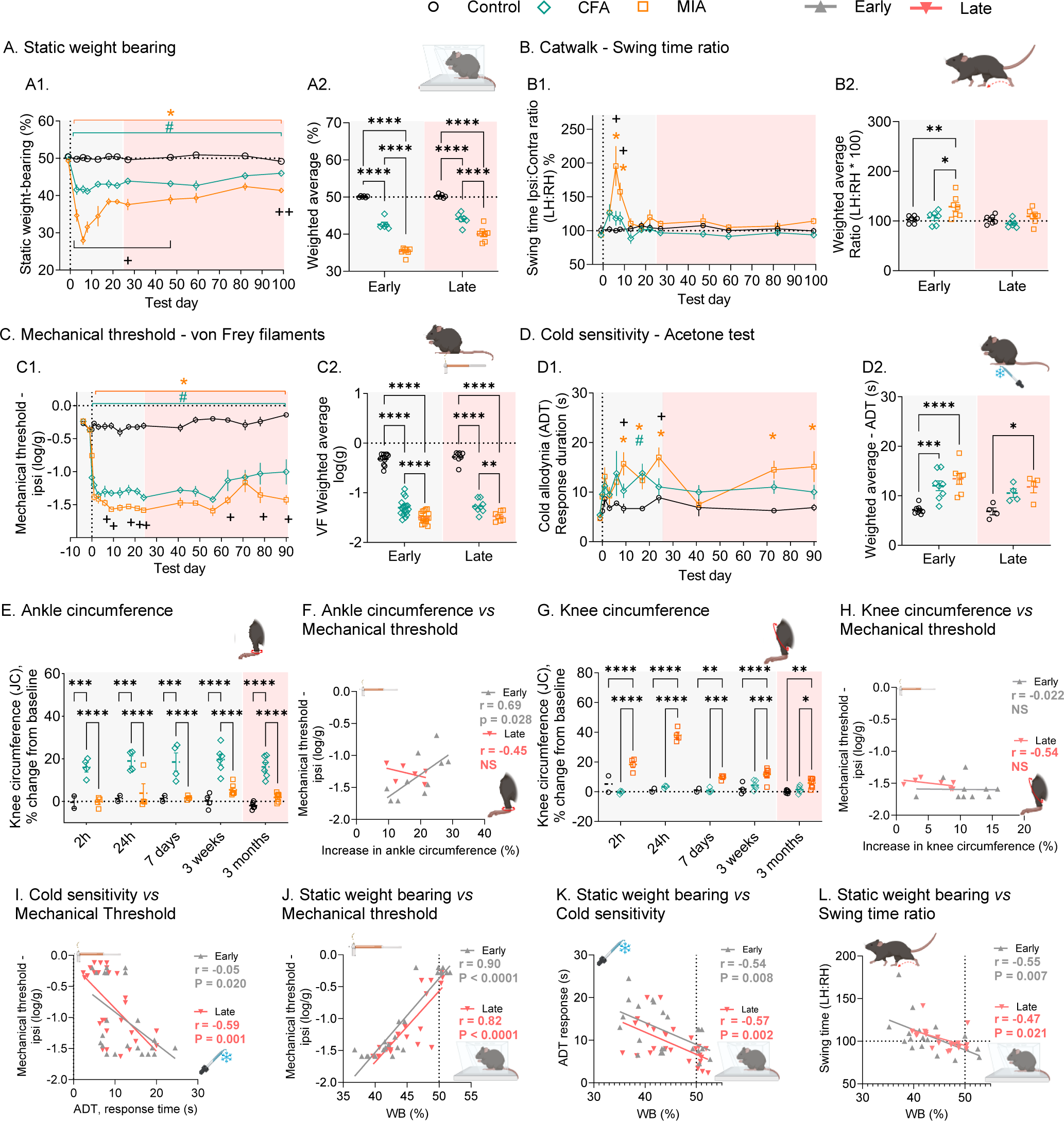
CFA and MIA induce persistent sensory and functional deficit that correlate for at least 3 months. **(A1,2)** Functional deficit following CFA and MIA injection was assessed using static weight bearing distribution (N=6/6/7). **(B1,2)** Functional deficit was also assessed by swing time ratio with the Catwalk gait analysis (N=6/6/7). **(C1,2)** Mechanical hypersensitivity was assessed using Von Frey filaments (N=4-18). **(D1,2)** Cold allodynia was determined using the acetone drop test (ADT) (N=4-8). **(E)** Ankle circumference was measured using calipers. **(F)** Ankle circumference in the early but not late phase of the disease states correlated with mechanical hypersensitivity (N=2-7). **(G)** Knee circumference was measured using calipers. **(H)** Knee circumference never correlated with mechanical hypersensitivity (N=2-7). **(I)** There was a correlation between mechanical threshold and cold sensitivity at both the early and late stage of the diseases states. **(J)** There was a correlation between mechanical threshold and static weight bearing distribution at both the early and late stage of the diseases states. **(K)** There was a correlation between static weight bearing distribution and cold sensitivity at both the early and late stage of the diseases states. **(L)** There was a correlation between swing time ratio and static weight bearing distribution at both the early and late stage of the diseases states. **Early**; Day 1-25, **Late**; Day 26-90. Data presented as individual time points and/or Mean ± S.E.M. Post-test in time-course figures (A1, B1, C1, D1): ^#^P<0.05, CFA *vs* control; *P<0.05, MIA *vs* control, ^+^P<0.05 CFA *vs* MIA, as determined using Tukey’s multiple comparison test. Full analysis-outcome in Supplementary Table S1. For scatterplot-figures (A2, B2, C2, D2, E, G), post-tests between injury-groups are displayed using connecting lines: *P<0.05, **P<0.01, ***P<0.001, ****P<0.0001, as determined using Tukey’s post-test.

To assess the sensory consequences of the injection of CFA into the ankle joint and MIA into the knee joint, we measured mechanical thresholds using von Frey monofilaments. A mixed effects model detected significant effect of both injury (P<0.0001) and time (P<0.0001) (Fig.1C1). At the earliest time point, *i.e.* 6 hours after injection, only the CFA-model produced significant allodynia, but from then on both the MIA and CFA-model produced significant mechanical allodynia compared to controls, from day 1 to day 90 after injury (P<0.05-0.0001). The MIA-group was significantly more sensitive than CFA on a few rare individual days (Fig.1C1). The significant difference between the CFA and MIA induced allodynia was also seen in the graph of weighted average across the two phases of the injury (Fig.1C2), where the MIA group displayed a significant lower mechanical threshold than CFA in both periods.

As cold allodynia is a significant hallmark of joint diseases, we also explored the response to acetone using the Acetone Drop Test (ADT). Both CFA and MIA induced long-lasting but highly dynamic cold-allodynia that was more prominent in MIA treated animals (Fig.1D1,2). There was a significant effect of both injury (P<0.0001) and time (P<0.0001), as determined using mixed effects model. Weighted averages analysis showed that both models displayed increased responses compared to control animals in the early phase, while in the late phase, only MIA induced significant cold allodynia compared with controls (Fig.1D2).

Animals were routinely monitored for body weight as well as ankle and knee joint swelling throughout the experiment. While animals seem to have been impacted by CFA within 24h of the injection, the bodyweight of both CFA and MIA animals was similar to that of control animals throughout the experiment (Fig.S1H). When we measured the swelling of the ankle and knee joint, we found that the CFA-induced swelling of the ankle was stable from 2h after the injection of CFA for up to 90 days (Fig.1E). Surprisingly, there was an inverse relationship between ankle swelling and mechanical threshold during the early phase in animals exposed to CFA-injection (Fig.1F). For the MIA model, following a steep increase, the swelling of the knee nearly returned to baseline by day 7 (Fig.1G) and there was no correlation between knee swelling and mechanical threshold for MIA animals (Fig.1H).

We finally explored whether there was any correlation between the sensory and functional outcomes measured and found a significant correlation at both the early and late phase of the joint pain development between mechanical thresholds and both cold allodynia (Fig.1I) and weight bearing (Fig.1J), between cold allodynia and weight bearing (Fig.1K), and between the measures of dynamic gait and static weight bearing (Fig.1L).

Overall, this data showed that both the MIA and CFA models induced a robust pain state that lasted at least 3 months, with small disparities in sensory outcomes but substantial differences in functional deficit, especially in the early stage of the disease states. We also found that there was a strong correlation between the parameters related to the sensory and functional outcome of joint pain, at both the early and the late stage of the disease states.

### CFA and MIA induce different patterns of neuronal and glia activation in the spinal cord

We next looked at molecular signaling in the superficial dorsal horn that could underlie the differences in sensory profile between the two models. To look at early signaling in the dorsal horn following CFA and MIA injection, we used immunohistochemistry to visualize c-Fos and CGRP expression. CFA and MIA induced similar pattern of c-Fos but distinct pattern of CGRP expression within 2h of injection (Fig.2A,B). Indeed, MIA induced greater expression of CGPR than seen in CFA and control animals, bilaterally (P<0.05) (Fig.2B).

**Figure 2.**
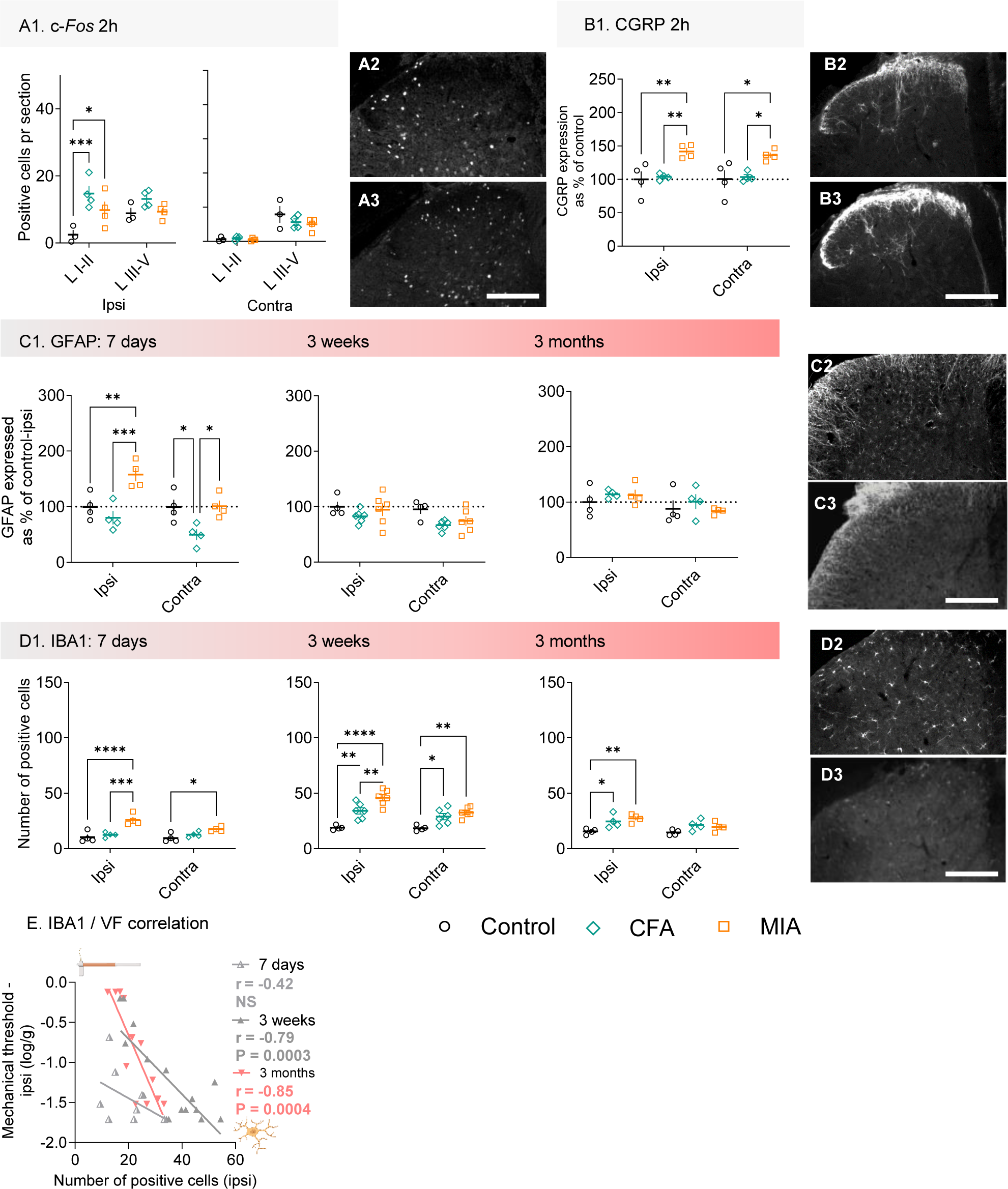
CFA and MIA induce different patterns of neuronal and glia activation in the spinal cord. **(A)** c-Fos expression was investigated 2h after the injection of CFA and MIA. Both CFA and MIA induced an upregulation of c-Fos in the ipsilateral superficial dorsal horn (**A1**). (**A2**) Representative picture of typical c-Fos expression in CFA mice in the ipsilateral superficial dorsal horn. (**A3**) Representative picture of typical cFos expression in MIA mice in the ipsilateral superficial dorsal horn. **(B)** CGRP expression was investigated 2h after the injection of CFA and MIA. MIA only induced a bilateral upregulation of CGRP (**B1**). (**B2**) Representative picture of typical CGRP expression in CFA mice in the ipsilateral superficial dorsal horn. (**B3**) Representative picture of typical CGRP expression in MIA mice in the ipsilateral superficial dorsal horn. **(C)** GFAP expression was investigated 7 days, 3 weeks and 3 months after the injection of CFA and MIA (**C1-3**). MIA only induced an upregulation of CGRP in the ipsilateral superficial dorsal horn (**C1**). (**C2**) Representative picture of typical GFAP expression in CFA mice in the ipsilateral superficial dorsal horn. (**C3**) Representative picture of typical CGRP expression in MIA mice in the ipsilateral superficial dorsal horn. **(D)** IBA1 expression was investigated 7 days, 3 weeks and 3 months after the injection of CFA and MIA (**D1-3**). MIA induced a bilateral upregulation of IBA1 (**D1**). (**D2**) Representative picture of typical IBA1 expression in CFA mice in the ipsilateral superficial dorsal horn. (**D3**) Representative picture of typical IBA1 expression in MIA mice in the ipsilateral superficial dorsal horn.; (**E**) IBA1 expression in the superficial dorsal horn correlates with the mechanical thresholds at 3 weeks and 3 months after initiation of the joint pain. Ipsi= ipslateral to injection; Contra = contralateral to injection. L I-II = laminae I and II; L III-V = laminae III to V. scale bar = 100µm; Post-test comparison of injury groups; N=3-6 per treatment group; *P<0.05, **P<0.01, ***P<0.001, ****P<0.0001, as determined using Tukey’s multiple comparison test.

We next looked at the activation of non-neuronal cells known to contribute to the maintenance of hypersensitive states and occurring later than neuronal activation. MIA induced a greater expression of the astrocytic marker GFAP at 7 days after injection (P<0.001, ipsi CFA vs MIA; P<0.05, contra CFA vs MIA) (Fig.2C). There was also a stronger up-regulation of the microglia marker IBA1 following MIA when compared to CFA (P<0.001, ipsi CFA vs MIA, 7 days and 3 weeks) (Fig.2D). IBA1 was upregulated after CFA only at 3 weeks and 3 months, but already from 7 days post-injury after MIA, and this MIA-induced up-regulation was bilateral at 7 days and 3 weeks. Finally, we looked at the correlation between these markers of nociceptive signalling and the mechanical hypersensitivity that develops after CFA and MIA and found that only IBA1 expression at 3 weeks and 3 months correlated with the degree of allodynia (Fig.2E).

### Activity and sleep patterns are more disrupted in the MIA than in the CFA model in the early phase of the persistent pain states

Because the differences in functional outcomes between the CFA and the MIA models were more prominent in the very early stages of the joint disease states, we decided to look at the activity and sleep patterns of a subset of mice in the first week after the injection of CFA and MIA. Mice were single-housed and home cage activity was tracked immediately following injury, using a system of passive infrared motion sensors [12] allowing to track activity and sleep for 7 consecutive days (Fig.3A). The injection of CFA and MIA, and therefore the start of the recordings, occurred just before the beginning of the dark phase at 19:00.

**Fig 3.**
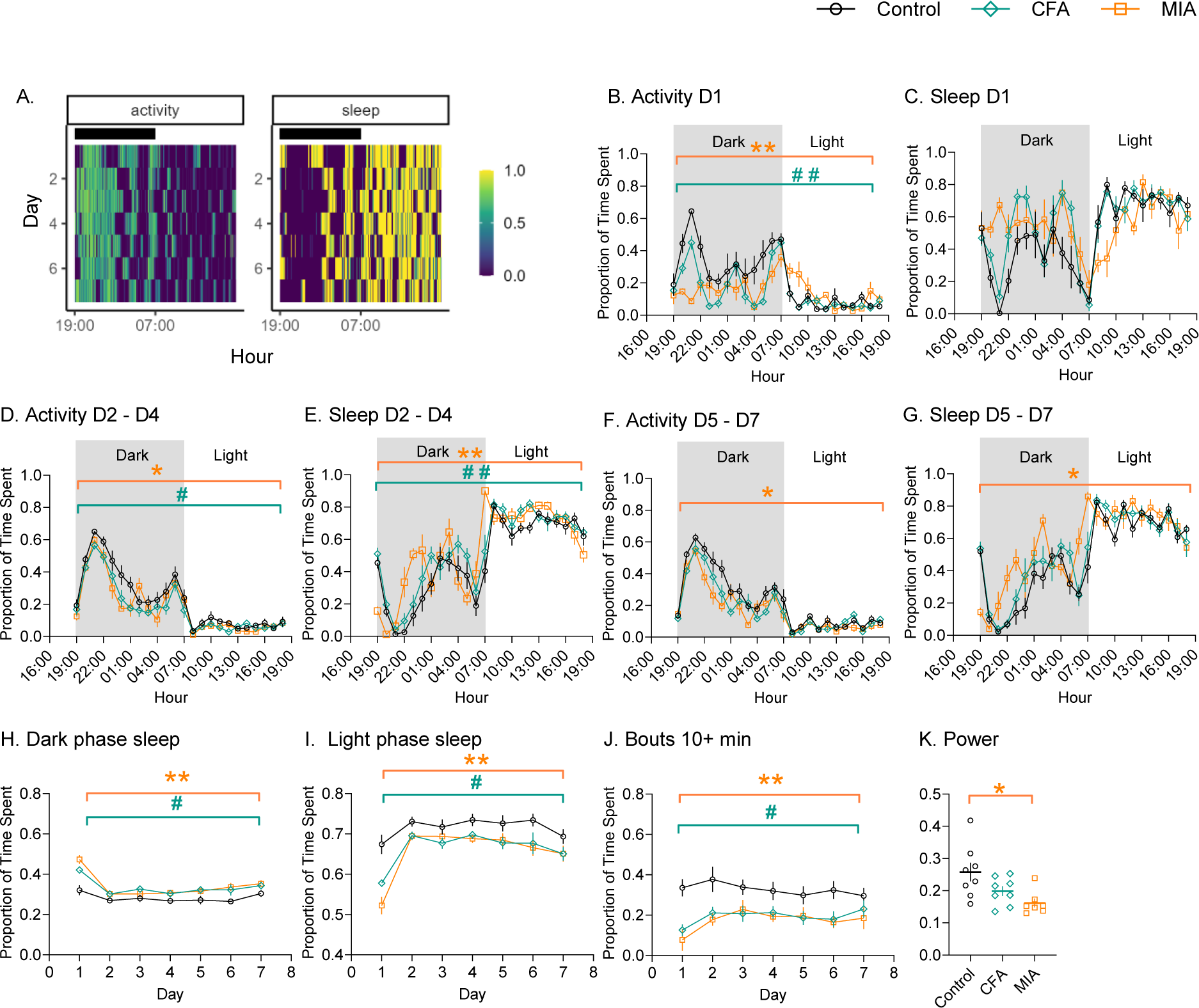
Sleep and activity patterns are more sensitive to MIA than CFA. (**A**) Representative activity and sleep patterns of a control mouse over 7 days of recordings, using 10 minute bins. (**B-G**) 24h activity/sleep plots using 1h bins. D1 = day 1 post CFA/MIA injection. D2-D4 = average of day 2 to day 4 post CFA/MIA injection. D5-D7 = average of day 5 to day 7 post CFA/MIA injection. (**H-J**) 7 day plots using 24h bins; (**H**) Proportion of sleep during the dark period. (**I**) Proportion of sleep during the light period. (**J**) Activity bouts of 10 min plus during the dark period. (**K**) A comparison of power of the 24-hour activity cycle calculated across 7 days. (**B-J**) Data shows mean ± S.E.M. (**K**) Single data points. N=8/8/7, Control/CFA/MIA; ^#^P<0.05, ^##^P<0.01, CFA *vs* control; *P<0.05, **P<0.05, MIA *vs* control, as determined using Tukey post-hoc analysis after repeated measures 2-way ANOVA.

Twenty four hour activity plots revealed that the activity and sleep patterns were most affected by CFA and MIA for the first 24h after the injections (Fig.3B,C), as animals recovered from the procedure. Overall, patterns of activity and sleep were similar across the 3 treatment groups from Day 2 after injection. Nonetheless, both CFA and MIA mice displayed reduced activity and increased sleep between Day 2 and Day 4 after injections (Fig.3D,E). While the injured animals started on the same activity trajectory as control animals, they seemed unable to sustain the same level of activity and instead showed increased sleep in the early hours of the dark phase. Between Day 5 and Day 7, only the MIA treated animals remained different from controls, while CFA treated mice behaved like control animals (Fig.3F,G).

Several summary statistics of circadian disruption [11] were calculated for individual animals across the 7 day period. Most of these statistics suggested that animals recovered rapidly following CFA and MIA injection, as there were little differences with control animals beyond day 1 after the procedure. Intra-daily variability measures how fragmented periods of activity are across a day [50]. Intra-daily variability was significantly higher in the day immediately following an injury, for both CFA and MIA mice, indicating more frequent changes between activity and rest, but quickly returned close to the same levels as the controls (Fig.S2A,B). Similarly, there was an immediate increase in light phase activity for injured animals and a decrease in dark-phase activity, but this was only observed on day 1 (Fig.S2C,D).

However, a number of statistics suggested longer lasting changes. Dark-phase sleep is a measure of the proportion of daily sleep that occurs when one would expect the animals to be active. Dark-phase sleep showed a significant increase in injured animals on the first day and remained higher than the control over a period of 7 days (Fig.3H). In contrast and as expected, light phase sleep was lower in injured animals compared to controls over the 7-day period (Fig.3I). The greatest effect of injury was observed when looking at the number of long bouts of activity that animals engaged with across the 7 days of observation. A bout is classed as a period of sustained activity above 10% of the mean activity level for the animal [14]. While short (0 to 1 minute in length) and medium (1 to 10 minutes) bouts of activity were no more common in injured than in control animals (Fig.S2E,F), longer bouts (greater than 10 minutes) were significantly less likely in CFA and MIA mice (Fig.2J). Finally, the Lomb-Scargle periodogram, similar to the chi-square periodogram [44], allows to evaluate how strong the activity-rest cycle is for different time periods (Fig.SG). Here, we saw a significant disruption in the 24-hour activity cycle for MIA animals only (Fig.3K).

Overall, these summary statistics suggested that there was not simply a shifting or spreading of activity into light periods, but that injured animals, in particular MIA animals, were less able to sustain activity during dark periods, requiring more frequent and prolonged rests.

### The MIA but not the CFA model induces robust negative affective behaviours

Last, we explored the affective response associated with joint pain. First, we looked at the duration of attentive response to a single application of three selected mechanical filaments (Low = 0.04g, Medium = 0.16g, High =1.0g). This affective response stabilised very rapidly after 10 days and was therefore only measured during the early phase of the pain states to reduce burden on the mice. For each of the three filaments, there was a significant effect of injury on the response duration when compared to controls and the MIA model induced stronger affective responses than the CFA model (Fig.4A, S3A,B,C). CFA animals actually rarely showed more affective behavior upon stimulation than control animals. Considering the large variation observed in the affective data collected, we decided to use an approach of individual behavioural profiling, developed to differentiate between “affected” and “exposed but unaffected” animals [3,45]. Affected animals were defined as responding one standard deviation or more from the average performance of the control group. Using these parameters, we found that only 40% of CFA animals displayed affective response to the application of Von Frey filament 0.04g *vs* 83% of MIA animals and 0% for control animals at 3 weeks post CFA and MIA injection (Fig.4A).

**Figure 4.**
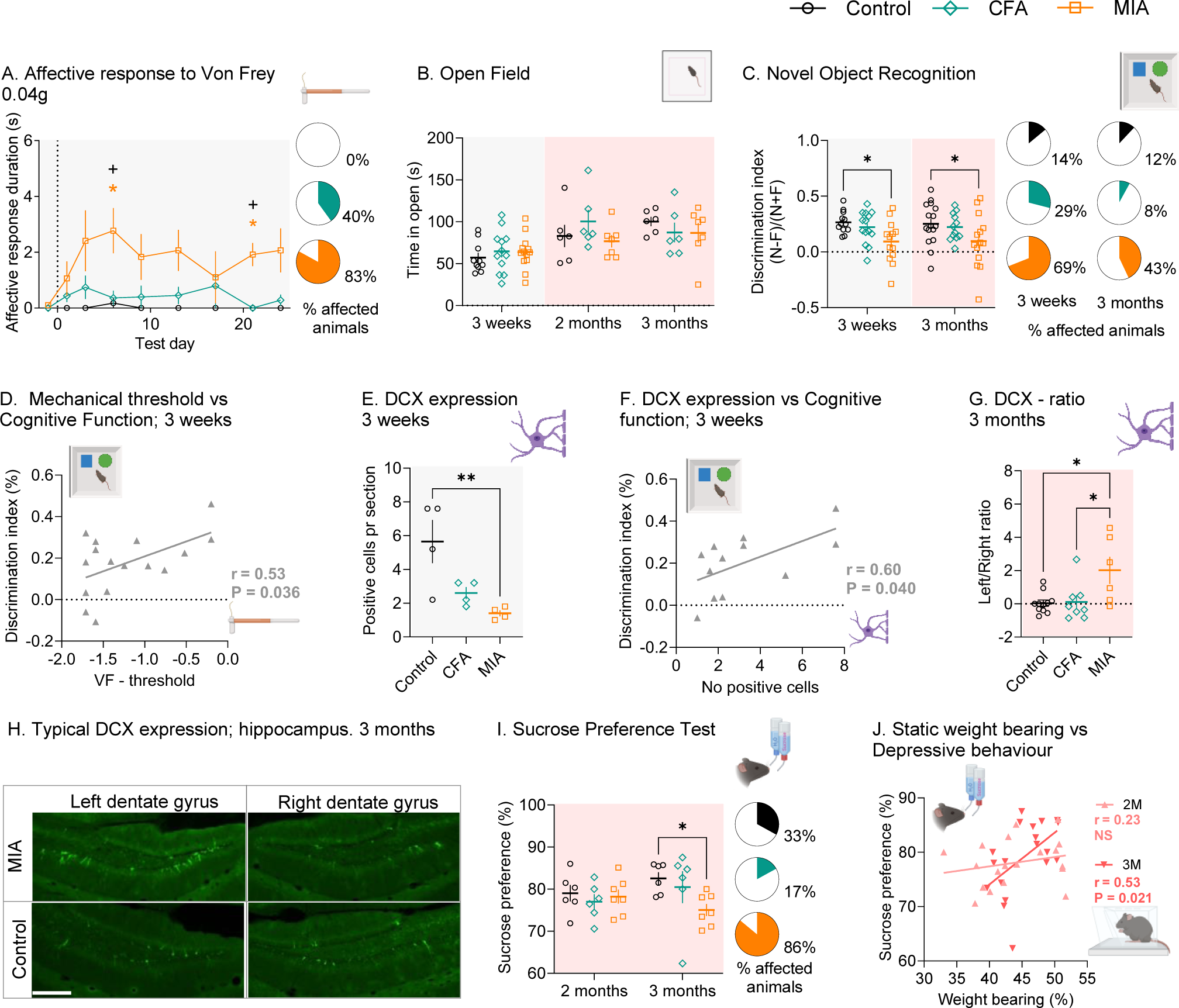
The MIA but not the CFA model induces robust negative affective behaviours. **(A)** The affective response to the application of the 0.04g Von Frey filament was recorded in seconds and affected animals were defined as animals that had a response greater than the control group average plus one standard deviation. N=4-10. (**B**) Anxiety-like behaviour was assessed using the time spent in the middle of the arena in the Open Field test at 3 weeks, 2 months and 3 months post MIA and CFA injection (N=6-13). (**C**) Novel Object Recognition was assessed at 3 weeks and 3 months post MIA and CFA injection. Discrimination index = (nose-interaction with novel / (nose interaction with novel + familiar))*100%. Affected animals were defined as animals that had a response lower than the control group average minus one standard deviation (N=12-16). (**D**) There was a correlation between mechanical threshold and cognitive deficit quantified by the discrimination index. (**E**) DCX expression in the hippocampus at 3 weeks post CFA and MIA injection was assessed using immunohistochemistry (N=4). (**F**) There was a correlation between the number of positive DCX cells in the hippocampus and cognitive deficit quantified by the discrimination index at 3 weeks after CFA and MIA injection. (**G**) DCX expression in the hippocampus at 3 months post CFA and MIA injection was assessed using immunohistochemistry. Here we present the ratio between the number of positive cells in the dentate gyrus of the left and right hippocampus (N=11/8/6). (**H**) Image representing the typical expression of DCX in the hippocampus in control and MIA animals, here in the left and right side at 3months after injury. White scale bar = 100µm. (**I**) The preference of sucrose against normal drinking water was measured in the Sucrose Preference Test to assess depressive like behaviour and affected animals were defined as animals that had a response lower than the control group average minus one standard deviation (N=6/6/7). (**J**) There was a correlation between static weight bearing deficit and depressive like behaviour at 3 months. Post-test in time-course figures (A); ^#^P<0.05, CFA *vs* control; *P<0.05, MIA *vs* control; ^+^P<0.05 CFA *vs* MIA, as determined using Tukey’s multiple comparison test. Full analysis-outcome in Supplementary Table S1. For scatter-plot figures post-tests between injury-groups are displayed using connecting lines *P<0.05, **P<0.01 as determined using Tukey’s post test.

Next, we explored the anxiety-like behaviour induced by CFA and MIA using the Open Field test and, unexpectedly, found little indication of anxiety-like behaviour in either joint pain model at 3 weeks, 2 months and 3 months following the induction of the pain state (Fig.4B). There was also no sign of differences in locomotion between the groups (Fig.S3D). As we were surprised by these results, we also used the Elevated Plus Maze to assess anxiety-like behavior in CFA and MIA treated mice, but again found no behavior suggesting anxiety (Fig.S3E).

However, when we next assessed cognitive function using the Novel Object Recognition (NOR) test, we observed significant cognitive deficit in the MIA model at 3 weeks and 3 months after the initiation of the pain state, but not in the CFA treated animals (Fig.4C, Fig.S3F,G). The decreased recognition of the novel object was seen when looking at the discrimination-index for the testing-phase alone (Fig.4C), but it was also clear that the MIA-injured animals displayed unequal amount of time at each of the two identical objects during the familiarisation phase, which was unmodified by the introduction of a novel object (Fig.S3F,G). Using individual behavioural profiling, we estimated that 29% of CFA animals displayed cognitive deficit *vs* 69% of MIA animals at 3 weeks (control, 14%), and 8% of CFA animals *vs* 43% of MIA animals at 3 months (control, 12%). We also found a correlation between the cognitive function and the mechanical sensitivity both measured at 3 weeks (Fig.4D). Importantly, when we looked at doublecortin (DCX) expressing neurons in the hippocampus that are important for learning [48], there was a significant decrease in DCX expression at 3 weeks in MIA animals (Fig.4E) and DCX expression level correlated with cognitive deficit (Fig.4F). We also looked at DCX expression at 3 months after CFA and MIA injection and while there were no obvious differences between the groups (Fig.S3H), there was an obvious laterality in DCX expression in the MIA group alone (Fig.4G,H).

We then looked at depressive-like behavior, using the sucrose preference test (SPT). Similarly, to the cognitive deficit assessment, we found that only MIA animals developed depressive-like behavior, as seen by reduced consumption of sucrose water. The development of depressive-like behaviour did not occur until 3 months after the initiation of the pain states (Fig.4I), at which time it correlated with the functional deficit estimated by static weight bearing (Fig.4J). Considering the within group variability, we again applied the individual behavioural profiling approach and, using control animal behaviour to identify affected *vs* non-affected animals, we found that 17% of CFA animals developed depressive-like behaviour *vs* 86% of MIA mice (control, 33%). All together these findings suggest that only MIA animals developed significant negative affective behavior.

### Not all pain related behaviours correlate

We wanted to explore how the different pain-related behaviors and selected molecular outcomes may correlate. Fig.5A,B summarize the correlations between outcome measures recorded for a same animal at 3 weeks (Fig.5A) and 3 months (Fig.5B), where the data was available. The correlation matrix clearly shows that the main sensory and functional outcomes (ADT, VF and WB) correlated well across models during both the early and late phase, and that changes in these outcome measures were also reflected in the molecular markers, specifically spinal IBA1 and hippocampal DCX. The emotional outcomes, on the other hand, did not consistently reflect the sensory outcomes. Moreover, despite WB and VF being strongly correlated at 3 weeks and 3 months, these outcomes were not equally correlated with the emotional outcomes. At 3 weeks, the cognitive deficits and affective response to Von Frey application correlated with mechanical allodynia (VF), while the depressive- and cognitive changes at 3 months correlated with the WB deficit. Similarly, the molecular markers showed significant correlation with NOR at 3 weeks, but not at 3 months.

**Figure 5:**
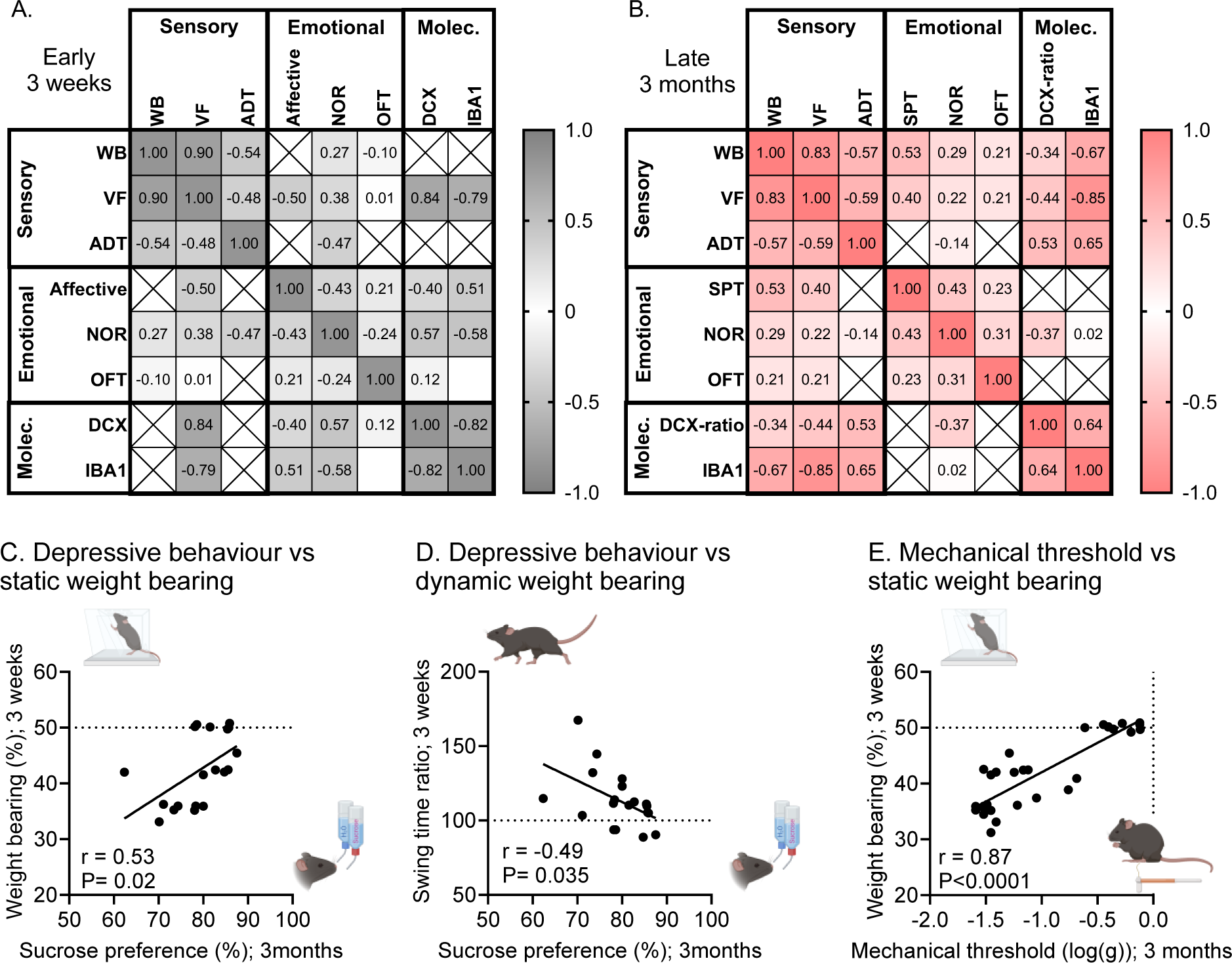
Early weight bearing deficit in joint disease can be used as a predictor of hypersensitivity and comorbid depressive-like behavior in late disease stage. Summary of correlations between outcome measures recorded on the same animal at the same time point at 3 weeks (**A**) and 3 months (**B**) after injury. Values displayed are r-values for Pearson r correlation analysis, estimating the strength of the correlations. The more intense color-coding signifies approaching the perfect fit at –1 or 1. (**C-E**) correlations between early weight bearing deficits at 3 weeks after injury and late outcomes at 3 months after injury. There was a significant correlation between (**C**) Depressive-like behavior at 3 months and static weight bearing deficit in the first 3 weeks after CFA and MIA injection; (**D**) Depressive-like behavior at 3 months and dynamic weight bearing in the first 3 weeks after CFA and MIA injection; and (**E**) Mechanical threshold at 3 months and static weight bearing deficit in the first 3 weeks after CFA and MIA injection. See full statistical analysis in supplementary table S1.

Finally, as predicting the development of significant comorbid affective disorders in patients with joint pain remains a significant clinical challenge, we asked whether early measures of functional deficit, which could be objectively evaluated in the clinics, may be used to predict the late development of comorbid affective disorders. Strikingly, we found that the early measures of both static and dynamic weight bearing (swing time) correlated with the depressive-like state as measured with the SPT at 3 months (Fig.5C,D). We then estimated the dependence of proportion of animals showing depressive behaviours at 3 months on weight bearing at 3 weeks, assigning animals to depressed vs non-depressed groups with the individual profiling approach. Using a generalised linear model with a logit link function, we found that the proportion of animals showing depressive behaviour decreased with more balanced weight bearing and that an estimate of 50% of animals with a weight bearing of 41.5% would show a depressive behavior (Fig.S4). Finally, there was also a very strong correlation between the early measures of static weight bearing and the late measures of allodynia (Fig.5E).

Our findings therefore suggest that the early functional deficit seen within the first 3 weeks of joint pain strongly correlated with the development of depressive-like behaviour seen 3 months after the onset of the disease, as well as the allodynia seen at this late time point.

## Discussion

Here we show that different models of joint pain may lead to overall similar sensory outcomes but strikingly different affective and molecular outcomes. Moreover, the very early stages of the joint pain states seem to be crucial for the long-term sensory and affective outcomes of the disease. Indeed, we found that the sensory and functional outcome of the MIA and CFA models differed more significantly in the early stage of the disease states and that the early changes in gait correlated with the likelihood of developing affective comorbidities such as depressive-like symptoms.

Our first unexpected finding was that while the CFA and MIA models were overall associated with similar sensory outcomes, only the MIA model showed significant affective comorbidities, such as cognitive-deficit, increased affective response to Von Frey application to the plantar surface, as well as depressive-like behaviour. These results were surprising, as we had hypothesised that the duration of the pain state would be a crucial factor for the development of affective comorbidities and both the CFA and MIA mice displayed significant allodynia for at least 3 months.

In disagreement with our findings, a recent study that monitored rats for up to 12 weeks after the injection of MIA in the knee joint failed to detect cognitive impairments [21]. However, the results obtained with rats indicated a reduced weight bearing deficit compared to our study and very little indication of allodynia, which is likely to explain the absence of cognitive deficit. Another recent study looking at mice with MIA injection in the knee joint supported our findings and demonstrated the presence of cognitive deficit as well as depressive-like behavior, but significantly earlier than we reported here [1]. Moreover, other experimental studies have previously shown changes in depressive-like behaviour and cognitive functions associated with hippocampus and pre-frontal cortex in rodent models of persistent pain [23,37,38], although timelines and occurrence of comorbid emotional changes in preclinical pain-models are very variable and at times contradicting [38,51].

Such drastic differences in affective outcomes between the 2 models highlights the importance of appropriately choosing pre-clinical models that resemble the human experience when investigating the mechanisms underlying long-term pain states and exploring novel therapeutic approaches. Moreover, while we observed very little variation in the sensory and functional outcome measures within each injured group, the variation was much greater when looking at affective behaviours. This is why we had to look at individual animal behavior, as others have done before in other research fields [3,45]. This approach allowed us to estimate that around 86% of MIA animals were presenting with depressive-like behavior and 57% with cognitive deficit, 3 months after disease onset. Osteo- and rheumatoid arthritis are well known to be accompanied by affective comorbidities with significant negative impact on patient well-being, but, similarly to what we observed in our mice, not all patients report affective comorbidities. About 17% of patients with rheumatoid arthritis have major depressive disorder [27] and crucially depression in patients with arthritis is associated with poor long-term outcomes, increased pain, fatigue and physical disability [27]. Therefore, it is paramount to get a better understanding of mechanisms underlying the development of the chronic pain states in those patients where comorbid depressive behavior must be addressed together with the pain that comes with joint disease. Future studies comparing animals that develop comorbid depression with non-affected animals should unveil new potential therapeutic targets for this particular subset of patients.

The MIA and CFA models used in this study showed significant differences in gait, all throughout the duration of the behavioural observations, but even more so in the early stages of the joint pain states. The MIA injury in particular produced compensatory mechanisms as seen by increased duration of the swing and length of the stride of the injured leg, and other output measures indicating a reduction of the load on the injured leg. Interestingly, it is also in the early days after MIA injection that mice displayed the strongest affective response to Von Frey filaments application to the plantar surface of the hindpaw, indicating that the early stage of the MIA disease was the most unpleasant to the animals. These observations were supported by our analysis of mice activity in their home cage, that clearly showed that mice that had received MIA had disrupted activity and sleeping patterns, as seen by their increased dark-phase sleep and reduced light-phase sleep, as well as their reduced engagement with long bouts (10min plus) of activity. Crucially, the measure of activity using the Open Field test did not show any differences in locomotion between the 3 treatment groups (control, CFA and MIA), indicating the importance of measuring mice activity in their home cage across few days, and not for a short period of time under experimental conditions, to get a better understanding of the impact of the intervention on their activity patterns. These results indicate that while locomotion may not be altered when observed in the context of a behavioural experiment, home cage observation might suggest otherwise.

Similar observations can be made when comparing the data obtained with the static weight bearing evaluation *vs* the Catwalk full gait analysis. Static weight bearing measures, which require the animals to sit still for a while and spread their weight between the 2 hindlegs, suggested clear and prolonged weight bearing deficit. However, in the Catwalk set-up, animals walk along the arena in a goal oriented manner and using 4 limbs, which lead to less signs of deficit with time. Indeed, all parameters reported showed similar peak deficits around one week after injury, with the dynamic Catwalk outcomes becoming progressively more subtle with time. Overall, the CFA and MIA models lead to different compensatory gait adaptation, with milder deficits from the CFA injury, which is likely to reflect the difference between an ankle and a knee injury [2].

According to our findings, the intense sensory, functional and affective changes in the early stages of the disease may be driving the long-term comorbidities such as depressive-like behaviour. While early weight bearing deficit did not correlate with the long-lasting cognitive deficit seen at 3 months, it correlated with the development of depressive like behavior and the intensity of the mechanical allodynia in the late stages of the joint diseases. Therefore, our findings suggest that early assessment of gait and pain related to load on the joint in patients with joint disease may be used as a predictor of long-term outcomes and early intervention may prove useful for better pain management and prevention of emotional comorbidities. However, one significant limitation of our work was the fast onset of the models used, as human joint pain does not develop in the short time frame observed with the CFA and MIA models. Indeed, common clinical symptoms including knee pain are very gradual in onset and pain worsens with time [25]. Therefore, comparing the time line of our “early” and “late” observations with those made in human patients is difficult. Moreover, our study was carried out in a restricted group of mice (8-week old at the onset of the disease, with similar weight and identical diet) while many factors are known to influence the outcome of joint diseases including age, sex and body mass index [31]. Sex differences in models of joint diseases have not often been studied, however a recent study suggest that microglia may be activated across sexes, but that glial inhibition may only reverse mechanical hypersensitivity in males [18]. Here, we found that glial activation in the dorsal horn was more prominent in the MIA than in the CFA model and that it correlated with the mechanical threshold at 3 weeks and 3 months. Whether this is also the case in female mice remains to be explored.

## Conclusions

Very few studies have investigated joint pain across different animal models to differentiate model specific behavioural and molecular outcomes *versus* those reflecting generalised chronic pain pathobiology [19,52]. Here, for the first time, we have identified correlation between early functional outcomes and late affective and sensory outcomes, particularly in the model of MIA that presented with stronger functional outcomes. Our results therefore suggest that early functional deficit may be a useful predictor of late comorbid depressive-like state and hypersensitivity, and that potential early intervention may prevent the later development of comorbid depression.

## Acknowledgements

We wish to thank Dr. Anna Learoyd for teaching of the use of the static weight bearing assay; Pia Jul for advice establishing the Novel Object Recognition assay; Dr. Maria Maiarù for teaching of model-induction and affective-motivational testing; and Dr. Lorenzo Fabrizi for advice on data analysis. This project was funded by a Versus Arthritis grant to SG. LM was supported by an Erasmus grant. SNP is funded by the Biotechnology and Biological Sciences Research Council (BB/X002357/1) and the National Centre for the Replacement, Refinement and Reduction of Animals in Research (NC/V000977/1).

