## Supplementary file for "Predicting hypersensitivity and comorbid depressive-like behavior in late stages of joint disease using early weight bearing deficit"

**Hestehave et al.**

### Supplementary data

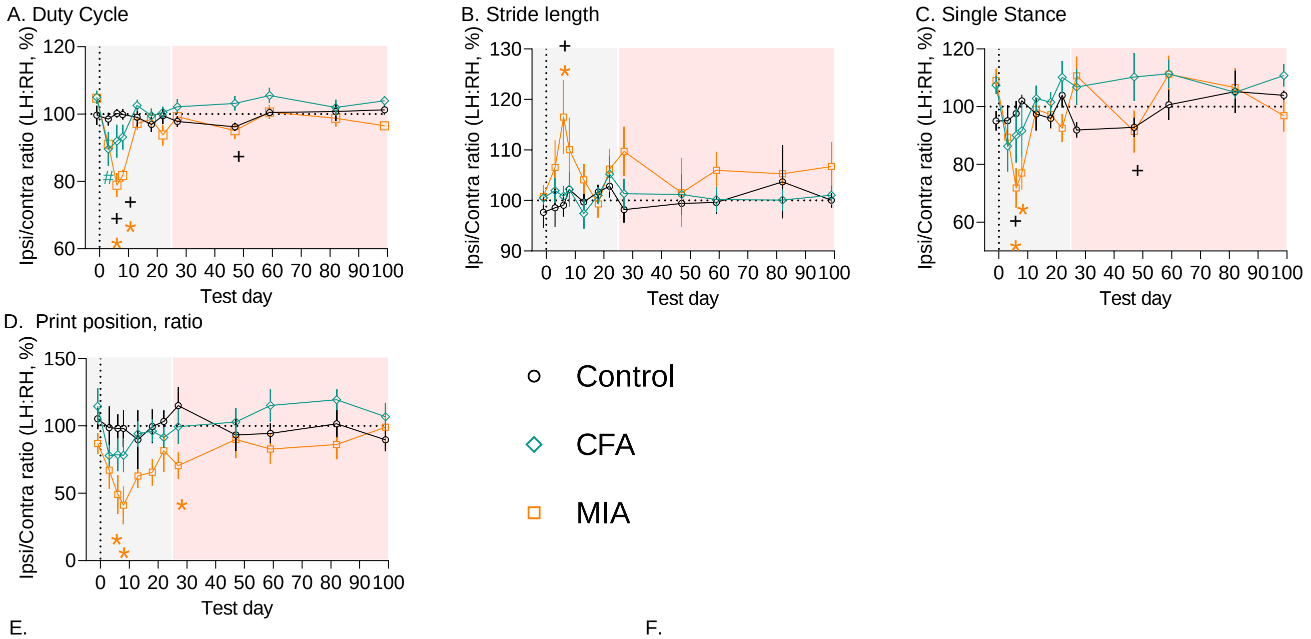


E.

| Peak Day 8 | Static WB | Single stance | Stride length | Duty cycle | Print position | Swing time |
|---|---|---|---|---|---|---|
| Static WB | 1.00 | 0.62 | -0.36 | 0.85 | 0.64 | -0.77 |
| Single stance | 0.62 | 1.00 | 0.22 | 0.81 | 0.34 | -0.60 |
| Stride length | -0.36 | 0.22 | 1.00 | -0.32 | -0.57 | 0.62 |
| Duty cycle | 0.85 | 0.81 | -0.32 | 1.00 | 0.73 | -0.86 |
| Print position | 0.64 | 0.34 | -0.57 | 0.73 | 1.00 | -0.68 |
| Swing time | -0.77 | -0.60 | 0.62 | -0.86 | -0.68 | 1.00 |

F.

| Early 3 weeks | Static WB | Single stance | Stride length | Duty cycle | Print position | Swing time |
|---|---|---|---|---|---|---|
| Static WB | 1.00 | 0.35 | -0.23 | 0.40 | 0.39 | -0.55 |
| Single stance | 0.35 | 1.00 | 0.14 | 0.72 | 0.14 | -0.43 |
| Stride length | -0.23 | 0.14 | 1.00 | -0.53 | -0.72 | 0.69 |
| Duty cycle | 0.40 | 0.72 | -0.53 | 1.00 | 0.68 | -0.81 |
| Print position | 0.39 | 0.14 | -0.72 | 0.68 | 1.00 | -0.70 |
| Swing time | -0.55 | -0.43 | 0.69 | -0.81 | -0.70 | 1.00 |

G.

| Late 3 months | Static WB | Single stance | Stride length | Duty cycle | Print position | Swing time |
|---|---|---|---|---|---|---|
| Static WB | 1.00 | 0.34 | -0.28 | 0.52 | -0.20 | -0.47 |
| Single stance | 0.34 | 1.00 | 0.46 | 0.84 | -0.44 | -0.14 |
| Stride length | -0.28 | 0.46 | 1.00 | 0.04 | -0.54 | 0.79 |
| Duty cycle | 0.52 | 0.84 | 0.04 | 1.00 | -0.15 | -0.53 |
| Print position | -0.20 | -0.44 | -0.54 | -0.15 | 1.00 | -0.35 |
| Swing time | -0.47 | -0.14 | 0.79 | -0.53 | -0.35 | 1.00 |

(WB: Static WB; Catwalk: Single stance, Stride length, Duty cycle, Print position, Swing time. Colour scale from -1.0 to 1.0.)

H.. Body weight gain

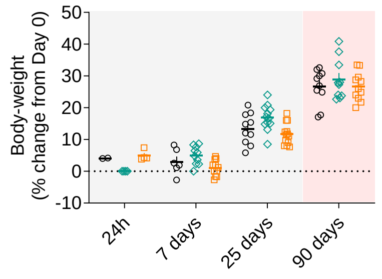

**Figure S1: MIA, but not CFA, induces prominent changes in dynamic gait, specifically in the early stages of the disease state**. **(A)** Duty Cycle, **(B)** Stride length, **(C)** Single stance and **(D)** Print Position were all affected by MIA injected in the knee joint. **(E-G)** Correlations between static weight bearing and catwalk outcome measures recorded in the same animal on the same day at Peak-Day 8, Early – 3 weeks and Late – 3months after injury. The r-values displayed signify the strength of the fit, as determined using Pearson r correlation analysis, and the higher intensity of the colors, the closer to the perfect fit at 1 or –1. **(H)** Body weights were monitored throughout the experiment and were no different from control animals. (**A-D**) Data shows mean ± S.E.M. Post-test in time-course figures (**A, B, C, D, H**); $^{\#}P<0.05$, CFA *vs* control; $^{*}P<0.05$, MIA *vs* control, $^{+}P<0.05$ CFA *vs* MIA, as determined using Tukey's multiple comparison test. Full analysis-outcome in Supplementary **Table S2**. N =6/6/7 for **A-D**. N=2-13 for **F**.

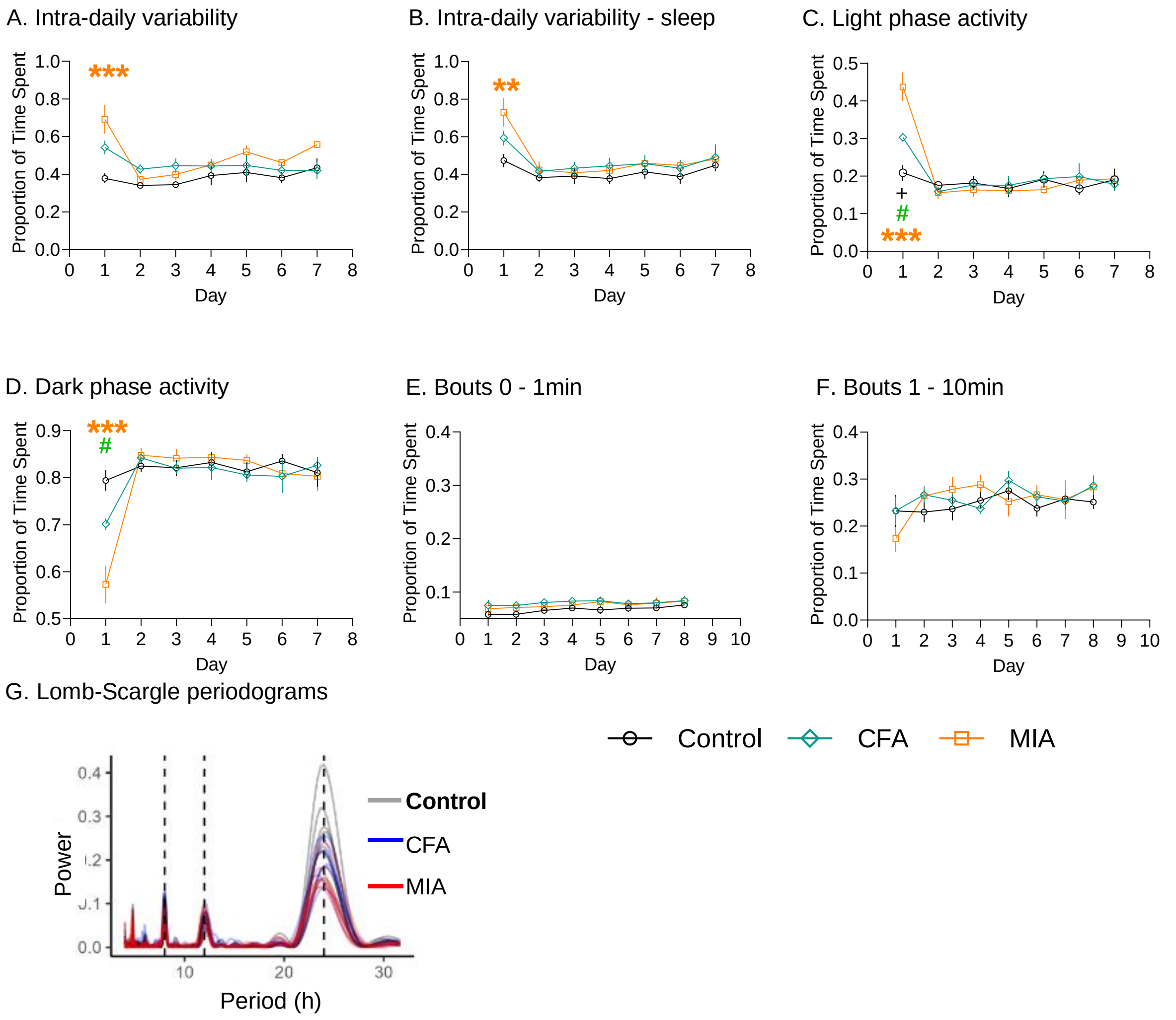


**Fig.S2: Sleep and activity patterns are more sensitive to MIA than CFA.** (**A-F**) 7-day plots using 24h bins; (**A**) Intra-daily variability. (**B**) Intra-daily variability sleep. (**C**) Proportion of activity during light period. (**D**) Proportion of activity during the dark period. (**E**) 0-1min bouts during the dark period recorded over 7 days. (**F**) 1-10min bouts during the dark period recorded over 7 days. (**G**) Lomb-Scargle periodograms for each mouse. Dashed lines mark 8, 12, and 24 hours. (**A-F**) Data shows mean ± S.E.M. N=8/8/7, Control/CFA/MIA; #P<0.05, CFA *vs* control; ***P<0.001, **P<0.05, MIA *vs* control, Univariate analysis at Day 1.

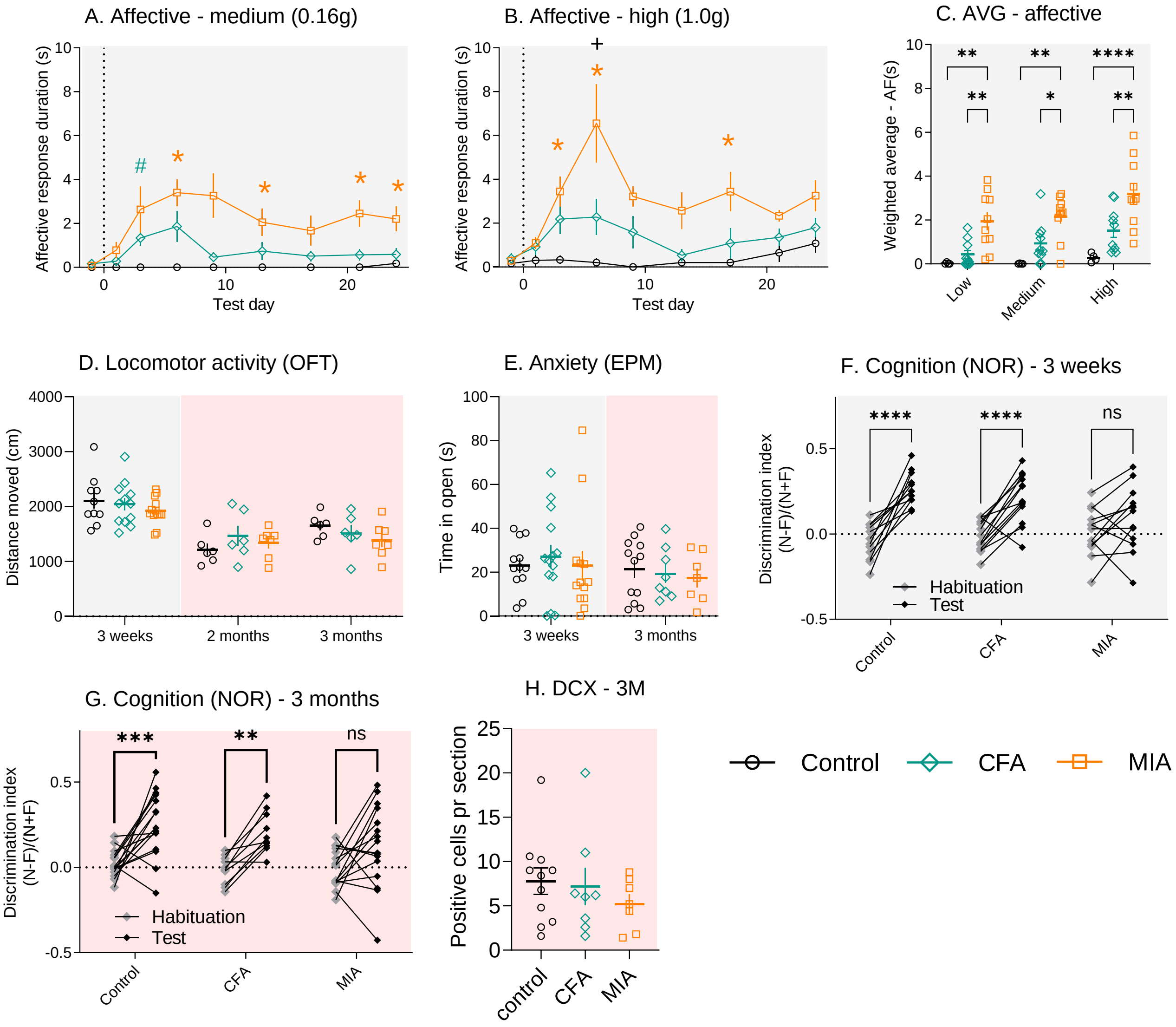


**Figure S3: The MIA but not the CFA model induces robust negative affective behaviours. (A, B)** The affective response to the application of the 0.16g and 1g Von Frey filaments was recorded in seconds (N=4-10). **(C)** Weighted average for the affective responses to all Von Frey filaments: low: 0.04g; Medium: 0.16g and High: 1g (N=4-10). **(D)** Locomotion was assessed using the distanced travelled in the Open Field test (N=6-13). **(E)** Anxiety like behaviour assessed using the amount of time exploring the open arms of the Elevated Plus Maze (N=7-14). **(F, G)** Discrimination index plots for the Novel Object Recognition test (N=12-16). **(H)** There was no difference in expression in DCX in the hippocampus at 3 months after CFA and MIA injections. (N=7-11). Post-test in time-course figures (A, B); $^{\#}$P<0.05, CFA *vs* control; *P<0.05, MIA *vs* control, $^{+}$P<0.05 CFA *vs* MIA, as determined using Tukey's multiple comparison test. Full analysis-outcome in Supplementary table S2. For (C,G,H) *P<0.05, **P<0.01, ***P<0.001, ****P<0.0001, as determined using appropriate post-test.

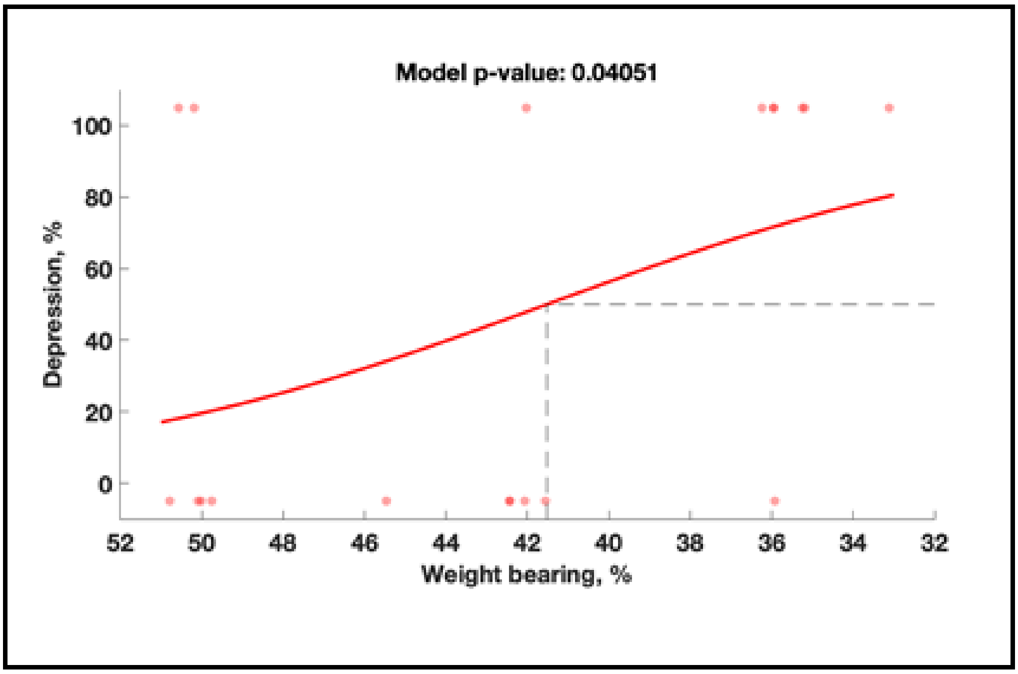


**Figure S4: Early weight bearing deficit in joint disease can be used to predict the development of depressive-like behavior in late disease stage.**

**Table S1. Statistical analysis table for main figures**

| Fig | Analysis[1] | F-values | Post test |
|---|---|---|---|
| **Fig 1 – sensory changes after injury** | | | |
| Fig 1A1 – Static Weightbearing | RM ANOVA, injury*time, | $F_{injury}$ (2,16) = 324.8, P<0.0001<br>$F_{time}$ (11,176) = 20.19, P<0.0001<br>$F_{interaction}$ (2,16) = 7.863, P<0.0001 | Tukey[2]<br>D3; ***, ##, +<br>D6; ****, ####, ++<br>D8; ****, ####, +++<br>D13; **, ####, +<br>D18; ****, ##, +<br>D22; ****, ####, ++<br>D27; ***, ###, +<br>D47; ***, ####, +<br>D59; ***, ##<br>D82; ***, ##,<br>D99; ****, #, ++ |
| Fig 1A2 – WB – weighted average | RM ANOVA, injury*phase. (early vs late considered as RM) | $F_{injury}$ (2,16) = 264.5, P<0.0001<br>$F_{phase}$ (1,16) = 25.56, P=0.0001<br>$F_{interaction}$ (2,16) = 10.99, P=0.001 | Tukey<br>Results displayed in figure. |
| Fig 1B1 – Swing time ratio | RM ANOVA, injury*time, | $F_{injury}$ (2,16) = 6.311, P=0.0095<br>$F_{time}$ (11,176) = 7.308, P<0.0001<br>$F_{interaction}$ (22,176) = 3.909, P<0.0001 | Tukey[2]<br>D6; ****, ++++<br>D8; ****, ++ |
| Fig 1B2 – Swing time – weighted average | RM ANOVA, injury*phase. (early vs late considered as RM) | $F_{injury}$ (2,16) = 5.690, P=0.0136<br>$F_{phase}$ (1,16) = 6.724, P=0.0196 | Tukey<br>Results displayed in figure. |
| Fig 1C1. – Von Frey – mechanical allodynia | Mixed Effects model[1], RM, injury*time | $F_{injury}$ (2,44) = 286.9, P<0.0001<br>$F_{time}$ (17,403) = 45.98, P<0.0001<br>$F_{interaction}$ (34,403) = 13.20, P<0.0001 | Tukey[2]<br>6H; ***, ####, +<br>D1 ; ****, ####, +<br>D3; ****, ####,<br>D6; ****, ####, +<br>D9; ****, ####, ++<br>D13; ****, ####,<br>D17; ****, ####, ++<br>D21; ****, ####, ++<br>D24; ****, ####, +<br>D41; ****, ####,<br>D48; ****, ####,<br>D56; ****, ####,<br>D63; ****, ####, +<br>D71; ****, ####,<br>D78; ****, ####, +<br>D90; ****, ####, ++ |
| Fig 1C2 – VF – weighted average | RM ANOVA, injury*phase. (Early vs late considered as RM) | $F_{injury}$ (2,64) = 479.9, P<0.0001 | Tukey<br>Results displayed in figure. |
| Fig 1D1 – Acetone Drop Test – cold allodynia | Mixed Effects model[1], RM, injury*time | $F_{injury}$ (2,20) = 18.46, p<0.0001<br>$F_{time}$ (10,144) = 6.36, p<0.0001 | Tukey[2]<br>D9; ****, +<br>D17; *, ###<br>D24; ***, +<br>D73; **<br>D90; ** |
| Fig 1D2 – ADT – weighted AVG. | Mixed Effects model, RM, injury*phase (early vs late considered as RM) | $F_{injury}$ (2, 29) = 17.92, P<0.0001 | Tukey<br>Results displayed in figure. |

| | | | |
|---|---|---|---|
| Fig 1E – ankle circumference | Two-way ANOVA, time*injury | $F_{injury}$ (2,50) = 90.06, P<0.0001 | Tukey Results displayed in figure. |
| Fig 1F – VF / ankle circumference correlation | Pearson r correlation Correlation of measures from early (Day 7+25) and late (3 months) | Early; r= 0.6857, P=0.0286.<br>Late; NS | |
| Fig 1G – knee circumference | Two-way ANOVA, time*injury | $F_{injury}$ (2,50) = 167.1, P<0.0001<br>$F_{time}$ (4,50) = 24.09, P<0.0001<br>$F_{interaction}$ (8.59) = 22.87, P<0.0001 | Tukey Results displayed in figure. |
| Fig 1H – VF / knee circumference correlation | Simple linear regression Correlation of measures from early (Day 7+25) and late (3 months) | NS | |
| Fig 1I – VF/ADT correlation | Pearsons r correlation, Single timepoint correlation from 3 weeks and 3 months. | 3 weeks/early; r=-.0480, P=0.020<br>3 months/Late; r= -0.594 P=0.001. | |
| Fig 1J - VF/WB correlation | Pearsons r correlation. Single timepoint correlation from 3 weeks and 3 months. | 3 weeks / early; r=0.898, P<0.0001.<br>3 months / late; r= 0.817, P<0.0001. | |
| Fig 1K - ADT/WB correlation | Pearsons r correlation Single timepoint correlation from 3 weeks and 3 months. | 3 weeks/early; r=-0.538, P=0.008<br>3 months/Late; r= -0.570, P=0.002. | |
| Fig 1L – Swing time / WB correlation | Pearsons r correlation Single timepoint correlation from 3 weeks and 3 months. | 3 weeks/early; r=-0.555, P=0.007<br>3 months/Late; r= -0.469 P=0.021. | |
| **Fig 2 – spinal molecular changes after injury** | | | |
| Fig 2A1. C-Fos 2h | Two-way RM ANOVA, area*injury (area = L II-II, L III-V x ipsi, contra treated as repeated measure) | $F_{injury}$(2,8) =8.2, P=0.012<br>$F_{area}$(3,24) =23.6, P<0.001<br>$F_{interaction}$(3,6) =4.2, P=0.029 | Tukey, Results displayed in figure. |
| Fig 2B1. CGRP – 2h | Two-way RM ANOVA, side*injury (ipsi-contra treated as repeated measure) | $F_{injury}$ (2,9) = 7.520, P=0.012 | Tukey Results displayed in figure. |
| Fig. 2C1; GFAP, I; 7 days, II; 3 weeks III; 3months | Two-way RM ANOVA, Side*injury (ipsi-contra treated as repeated measure) | D7;$F_{injury}$ (2,9) = 8.103, P=0.0097<br>$F_{side}$ (1,9) = 138.3, P<0.0001<br>$F_{interaction}$ (2,9) = 42.31, P<0.0001<br>3W; $F_{side}$ (1,13) = 11.90, P=0.0043<br>3M; $F_{side}$ (1,9) = 12.67, P=0.0061 | Tukey Results displayed in figure. |
| Fig. 2D1; IBA1, I; 7 days, II; 3 weeks III; 3months | Two-way RM ANOVA, Side*injury (ipsi-contra treated as repeated measure) | D7;$F_{injury}$ (2,9) = 14.88, P=0.0014<br>$F_{side}$ (1,9) = 7.590, P=0.0223<br>$F_{interaction}$ (2,9) = 6.417, P=0.0185<br>3W;$F_{injury}$ (2,13) = 15.75, P=0.0003<br>$F_{side}$ (1,13) = 114,7, P<0.0001<br>$F_{interaction}$ (2,13) = 40.27, P<0.0001<br>3M;$F_{injury}$ (2,9) = 5.707, P=0.0251<br>$F_{side}$ (1,9) = 22.30, P=0.0011<br>$F_{interaction}$ (2,9) = 5.124, P=0.0324 | Tukey Results displayed in figure. |
| Fig. 2E. IBA1/VF correlation at 7 days, 3 weeks, 3 months, | Pearson r Correlation of measures from day 7, 3 weeks, 3 months | 7 days; r=-0.4194 P=0.30 (NS)<br>3 weeks/early; r=-0.7905, P=0.0003<br>3 months/Late; r= -0.8510, P=0.0004. | |

| | | | |
|---|---|---|---|
| **Fig 3. Sleep and activity pattern after injury** | | | |
| Fig 3B. Activity D1 | Two-way RM ANOVA, time*injury | $F_{time}(23,460)$ =16.9, P<0.001<br>$F_{injury}(2,20)$ =9.9, P<001<br>$F_{interactions}(46,460)$ =3.7, P<0.001 | Tukey for Injury Results displayed in figure. |
| Fig 3C. Sleep D1 | Two-way RM ANOVA, time*injury | $F_{time}(23,460)$ =13.4, P<0.001<br>$F_{injury}(2,20)$ =3.3, P=0.059<br>$F_{interactions}(46,460)$ =3.1, P<0.001 | |
| Fig 3D. Activity D2-D4 | Two-way RM ANOVA, time*injury | $F_{time}(23,460)$ =81, P<0.001<br>$F_{injury}(2,20)$ =5.1, P=0.017<br>$F_{interactions}(46,460)$ =1.5, P=0.128 | Tukey for Injury Results displayed in figure. |
| Fig 3E. Sleep D2-D4 | Two-way RM ANOVA, time*injury | $F_{time}(23,460)$ =57.9, P<0.001<br>$F_{injury}(2,20)$ =8.2, P=0.002<br>$F_{interactions}(46,460)$ =4.1, P<0.001 | Tukey for Injury Results displayed in figure. |
| Fig 3F. Activity D5-D7 | Two-way RM ANOVA, time*injury | $F_{time}(23,460)$ =67.6, P<0.001<br>$F_{injury}(2,20)$ =5.1, P=0.016<br>$F_{interactions}(46,460)$ =1.8, P=0.029 | Tukey for Injury Results displayed in figure. |
| Fig 3G: Sleep D5-D7 | Two-way RM ANOVA, time*injury | $F_{time}(23,460)$ =46.5, P<0.001<br>$F_{injury}(2,20)$ =4.6, P=0.023<br>$F_{interactions}(46,460)$ =3.5, P<0.001 | Tukey for Injury Results displayed in figure. |
| Fig 3H: Dark phase sleep | Two-way RM ANOVA, time*injury | $F_{time}(6,120)$ =27.7, P<0.001<br>$F_{injury}(2,20)$ =8.1, P=0.003<br>$F_{interactions}(12,120)$ =3.5, P=0.012 | Tukey for Injury Results displayed in figure. |
| Fig 3I: Light phase sleep | Two-way RM ANOVA, time*injury | $F_{time}(6,120)$ =28.7, P<0.001<br>$F_{injury}(2,20)$ =8.1, P=0.003<br>$F_{interactions}(12,120)$ =2.6, P=0.017 | Tukey for Injury Results displayed in figure. |
| Fig 3J: Bouts 10+ min | Two-way RM ANOVA, time*injury | $F_{time}(6,120)$ =2.46, P=0.076 (NS)<br>$F_{injury}(2,20)$ =1.2, P=0.309<br>$F_{interactions}(12,120)$ =7.1, P=0.005 | Tukey for Injury Results displayed in figure. |
| Fig 3K. Power | Univariate analysis | $F_{injury}(2,20)$ =5.2, P=0.015 | Tukey for Injury Results displayed in figure. |
| **Fig 4 – emotional/affective comorbidities** | | | |
| Fig 4A – Affective responding – low (0.04g) | Mixed Effects model[1], RM, injury*time | $F_{injury}$ (2,21) = 15.74, P<0.0001 | Tukey[2]<br>D6; *, +<br>D21; *, + |
| Fig 4B – Anxiety (OFT) | Two-way ANOVA, time*injury | $F_{time}$ (2, 64) = 11.26, P<0.0001<br>$F_{injury}$ = NS | - |
| Fig 4C – NOR disc index. | Two-way ANOVA, injury*time (not RM) | $F_{injury}$ (2, 75) = 6.783, P=0.002 | Tukey Results displayed in figure. |
| Fig 4D – NOR/VF correlation, 3W | Pearson r correlation | r=0.5279, P=0.0356 | |
| Fig 4E – DCX 3 weeks | One way ANOVA, injury | $F_{injury}$ (2, 9) = 7.990, P=0.0101 | Tukey Results displayed in figure. |
| Fig 4F – NOR/DCX correlation, 3W | Pearson r correlation | r=0.5992, P=0.0391 | |
| Fig 4G – DCX 3M, left/right ratio | One-way ANOVA | $F_{injury}$ (2, 22) = 5.820, P=0.0094 | Tukey Results displayed in figure. |
| Fig 4I – Anhedonia (SPT) | Two-way RM ANOVA, Time*injury | $F_{time}$ (1, 64) =0.7773, P=0.3910 = NS<br>$F_{injury}$ (2, 16) = 1.513, P=0.2501 = NS<br>$F_{time*injury}$ (2, 16) = 2.515, P=0.1122 (NS) | Tukey Results displayed in figure. |

| | | |
|---|---|---|
| Fig 4J – SPT/WB correlation | Pearson r correlation at 2 and 3 months | 2 Months; P=0.35 (NS)<br>3 Months; r=0.5264, P=0.0206 |
| **Fig 5 – correlation matrix** | | |
| Fig 5A. 3 weeks / Early correlation matrix | Pearson r correlation (only correlations that are significant, or below P=0.1 are presented here) | WB/VF; r=0.898, P<0.0001.<br>WB/ADT; r=-0.538, P=0.008.<br>VF/ADT; r=-0.480, P=0.020<br>VF / affective-low; r=-0.500, P=0.049.<br>VF/NOR; r=0.385, P=0.016.<br>VF/DCX; r=0.837, P=0.0007.<br>VF/IBA-1; r=-0.791, P=0.0003.<br>ADT/NOR; r=-0.470, P=0.0236.<br>NOR/DCX; r=0.569, P=0.0395.<br>NOR/IBA-1; r=-0.584, P=0.0175.<br>DCX/IBA-1; r=-0.815, P=0.0012.<br>IBA-1/affective-low; r=0.531, P=0.042. |
| Fig 5B. 3 months / Late correlation matrix | Pearson r correlation (only correlations that are significant, or below P=0.1 are presented here) | WB/VF; r= 0.833, P<0.0001.<br>WB/ADT; r= -0.570, P=0.0019.<br>WB/NOR; r=0.288, P=0.065 (NS).<br>WB/SPT; r=0.526, P=0.0206.<br>WB/IBA-1; r= -0.668, P=0.0176<br>VF/ADT; r= -0.594, P=0.001.<br>VF/SPT; r=0.403, P=0.0869 (NS)<br>VF/DCX-ratio; r=-0.442, P=0.0446<br>VF/IBA-1; r= -0.851, P=0.0004<br>ADT/DCX-ratio; r= 0.526, P=0.0144<br>ADT/IBA-1; r= 0.648, P=0.0227<br>SPT/NOR; r=0.426, P=0.0690 (NS)<br>NOR/DCX-ratio; r=-0.367, P=0.1013 (NS)<br>DCX-ratio/IBA-1; r=0.638, P=0.0471. |
| Fig 5C. Depressive behaviour vs static WB | Pearson r correlation | r=0.5314, P=0.0192 |
| Fig 5D. Depressive behaviour vs dynamic weight bearing (swing ratio) | Pearson r correlation | r=-0.4892, P=0.0335 |
| Fig 5E. Mechanical threshold vs static weight bearing | Pearson r correlation | r=0.8673, P<0.0001 |

[1] Mixed effects model due to different duration of experimental cohorts, meaning missing values. All cohorts though included equal numbers of animals from the different groups. Weighted average was calculated based on the period that the animals in question was tested.

[2]Post-tests for time-course figures; MIA vs control; *; *P<0.05, **P<0.01, ***P<0.001. ****P<0.0001. CFA vs control; #; #P<0.05, ##P<0.01, ###P<0.001. ####P<0.0001. CFA vs MIA; +; +P<0.05, ++P<0.01, +++P<0.001. ++++P<0.0001.

RM= Repeated Measures

Additional stats: 3weeks AVG vs 3 month outcome;

- Early-WB / Late- DCX-ratio; r=-0.4794, P=0.0279.
- Early-WB / Late-IBA-1; r=-0.7313, P=0.0069.
- Early-ADT/ Late-IBA-1; r=0.6705, P=0.0170.

### Table S2. Statistical analysis table for main figures

| Fig | Analysis | F-values | Post test |
|---|---|---|---|
| **Fig S1 – catwalk and bodyweight** | | | |
| Fig S1A – Duty Cycle | RM ANOVA, injury*time | $F_{injury}$ (2,16) = 4.904, P=0.0218<br>$F_{time}$ (11,176) = 11.15, P<0.0001<br>$F_{interaction}$ (22,176) = 4.129, P<0.0001 | Tukey[2]<br>D3; #<br>D6; ****, +++<br>D8; ****, ++<br>D47; + |
| Fig S1B – Stride Length | RM ANOVA, injury*time | $F_{injury}$ (2,16) = 2.853, P=0.0872 (NS) | Tukey[2]<br>D6; **, + |
| Fig S1C – Single Stance | RM ANOVA, injury*time, | $F_{injury}$ (2,16) = 1.655, P=0.2222 (NS)<br>$F_{time}$ (11,176) = 5.450, P<0.0001<br>$F_{interaction}$ (22,176) = 2.414, P=0.0008 | Tukey[2]<br>D6; **, +<br>D8; **<br>D27; *<br>D47; + |
| Fig S1D – Print position | RM ANOVA, injury*time | $F_{injury}$ (2,16) = 3.155, P=0.07 (NS)<br>$F_{time}$ (11,176) = 4.045, P<0.0001<br>$F_{interaction}$ (22,176) = 1.585, P=0.054 (NS) | Tukey[2]<br>D6; **<br>D8; **<br>D27; * |
| Fig S1H – body weight gain | RM ANOVA, injury*time, | $F_{injury}$ (2,91) = 1.151, P=0.3210 (NS)<br>$F_{time}$ (2,91) = 183.5, P<0.0001<br>$F_{interaction}$ (6,91) = 1.679, P=0.1352(NS) | |
| Fig S1E. Peak – Day 8 correlation matrix | Pearson r correlation matrix (only correlations that are significant, or below P=0.1 are presented) | WB/Swing; r=-0.769, P=0.0001<br>WB/Duty cycle; r=0.846, P<0.0001<br>WB/Stride length; r=-0.362, P=0.064 (NS)<br>WB/Single stance; r=0.617, P=0.002<br>WB/Print position; r=0.641, P=0.002<br>Single stance/print position; r= 0.338, p=0.079 (NS)<br>Single stance / swing time; r= -0.599, P=0.003<br>Single stance / duty cycle; r=0.812, P<0.0001<br>Stride length / duty cycle; r=-0.322 , P=0.089 (NS)<br>Stride length / print position; r= -0.571, P=0.005<br>Stride length / swing time; r= 0.624, P=0.002<br>Duty cycle / print position; r= 0.727, P<0.0001<br>Duty cycle / swing time; r=-0.864 . P<0.0001<br>Print position / swing time; r= -0.680, P=0.0001 | |
| Fig S1F. Early – 3W correlation matrix | Pearson r correlation matrix (only correlations that are significant, or below P=0.1 are presented) | WB/Swing; r=-0.555, P=0.007<br>WB/Duty cycle; r=0.404, P=0.043<br>WB/Single stance; r=0.354, P=0.0.069(NS)<br>WB/Print position; r=0.393, P=0.048<br>Single stance / duty cycle; r= 0.720, P<0.0001<br>Single stance / swing time; r= -0.435, P=0.031<br>Stride length / duty cycle; r= -0.533, P=0.009<br>Stride length / print position; r= -0.722, P<0.0001<br>Stride length / swing time; r= 0.687, P=0.001<br>Duty cycle / print position; r= 0.679, P=0.001<br>Duty cycle / swing time; r= -0.808, P<0.0001<br>Print position / swing time; r= -0.701, P<0.0001 | |
| Fig S1G. Late – 3M correlation matrix | Pearson r correlation matrix (only correlations that are significant, or below P=0.1 are presented) | WB/Swing; r=-0.469, P=0.021<br>WB/Duty cycle; r=0.519, P=0.011<br>WB/Single stance; r=0.338, P=0.078 (NS)<br>Single stance / stride length; r= 0.463, P=0.023<br>Single stance / duty cycle; r= 0.843, P<0.0001<br>Single stance / print position; r= -0.438, P=0.030<br>Stride length / print position; r= -0.543, P=0.008<br>Stride length / swing time; r= 0.790, P<0.0001<br>Duty cycle / swing time; r= -530, P=0.01<br>Print position / swing time; r= -0.352, P=0.07 (NS) | |

| Fig S2 – additional sleep | | | |
|---|---|---|---|
| Fig S2A. Intra – daily variability | Two-way RM ANOVA, time*injury N.B. here only D1 to D6 as missing data for D7 | $F_{time}(5,100)$ =16.9, P<0.001<br>$F_{injury}(2,20)$ =5.0, P=0.017<br>$F_{interactions}(10,100)$ =3.3, P=0.008 | Tukey for Injury at D1 Results displayed in figure. |
| Fig S2B. Intra – daily variability sleep | Two-way RM ANOVA, time*injury N.B. here only D1 to D6 as missing data for D7 | $F_{time}(5,100)$ =24.5, P<0.001<br>$F_{injury}(2,20)$ =1.5, P=0.246<br>$F_{interactions}(10,100)$=3.0, P=0.007 | Univariate at D1 displayed in figure. |
| Fig S2C. Light-phase activity - | Two-way RM ANOVA, time*injury | $F_{time}(6,120)$ =31.6, P<0.001<br>$F_{injury}(2,20)$ =0.852, P=0.442<br>$F_{interactions}(12,120)$ =7.6, P<0.001 | Univariate at D1 displayed in figure. |
| Fig S2D. Dark-phase activity | Two-way RM ANOVA, time*injury | $F_{time}(6,120)$ =27.3, P<0.001<br>$F_{injury}(2,20)$ =0.84, P=0.447<br>$F_{interactions}(12,120)$ =6.8, P<0.001 | Univariate at D1 displayed in figure. |
| Fig S2E. Bouts 0-1min | Two-way RM ANOVA, time*injury | $F_{time}(6,120)$ =1.6, P=0.147<br>$F_{injury}(2,20)$ =1.9, P=0.176<br>$F_{interactions}(12,120)$ =0.2, P=0.98 | |
| Fig S2F. Bouts 1-10 min | Two-way RM ANOVA, time*injury | $F_{time}(6,120)$ =3.2, P=0.026<br>$F_{injury}(2,20)$ =0.156, P=0.857<br>$F_{interactions}(12,120)$ =1.52, P=0.181 | |
| Fig S3 – emotional supplementary | | | |
| Fig S3A. Affective medium (0.16g) | Mixed Effects model[1], RM, injury*time. | $F_{injury}$ (2,21) = 23.47, P<0.0001 | Tukey[2]<br>D3; #<br>D6; ***<br>D13; *<br>D21; *<br>D24; * |
| Fig S3B. Affective high (1g) | Mixed Effects model[1], RM, injury*time. | $F_{injury}$ (2,21) = 11.40, P=0.0004<br>$F_{time}$ (8,128) = 3.457, P=0.0012<br>$F_{interaction}$ (8,128) = 3.457, P=0.0012 | Tukey<br>D3; *<br>D6; ****, ++++<br>D17; * |
| Fig S3C – affective weighted average | RM ANOVA, force*injury. (different filament forces considered as RM) | $F_{force}$ (2, 42) = 9.220, P=0.0005<br>$F_{injury}$ (2, 21) = 12.58, P=0.0003<br>$F_{subject}$ (21, 42) = 5.171, P<0.0001 | Tukey[2] Results displayed in figure |
| Fig S3D. Open field – distance | Two-way ANOVA, time*injury | $F_{time}$ (2,64) = 28.07, P<0.0001 | |
| Fig S3E. Elevated Plus Maze | Two-way ANOVA, time*injury | NS | |
| Fig S3F. Novel Object Recognition, 3W | RM ANOVA, injury*test-phase. | $F_{test\text{-}phase}$ (1,36) = 63.94, P<0.0001<br>$F_{interaction}$ (2,36) = 6.534, P=0.0038 | Sidak, Results displayed in figure. |
| Fig S3G. Novel Object Recognition, 3M | RM ANOVA, injury*test-phase. | $F_{test\text{-}phase}$ (1,39) = 31.70, P<0.0001<br>$F_{injury}$ (2,39) = 2.087, P=0.1376 (NS) | Sidak, Results displayed in figure. |
| Fig 3E – DCX 3M. | One-way ANOVA | P= 0.55 (NS) | |

[1] Mixed effects model due to different duration of experimental cohorts, meaning missing values. All cohorts though included equal numbers of animals from the different groups. Weighted average was calculated based on the period that the animals in question was tested.

[2]Post-tests for time-course figures; MIA vs control; *; *P<0.05, **P<0.01, ***P<0.001. ****P<0.0001. CFA vs control; #; #P<0.05, ##P<0.01, ###P<0.001. ####P<0.0001. CFA vs MIA; +; +P<0.05, ++P<0.01, +++P<0.001. ++++P<0.0001.
